# Biodiversity in mountain soils above the treeline

**DOI:** 10.1101/2023.12.22.569885

**Authors:** Nadine Praeg, Michael Steinwandter, Davnah Urbach, Mark A. Snethlage, Rodrigo P. Alves, Martha E. Apple, Andrea J. Britton, Estelle P. Bruni, Ting-Wen Chen, Kenneth Dumack, Fernando Fernandez-Mendoza, Michele Freppaz, Beat Frey, Nathalie Fromin, Stefan Geisen, Martin Grube, Elia Guariento, Antoine Guisan, Qiao-Qiao Ji, Juan J. Jiménez, Stefanie Maier, Lucie A. Malard, Maria A. Minor, Cowan C. Mc Lean, Edward A. D. Mitchell, Thomas Peham, Roberto Pizzolotto, Andy F. S. Taylor, Philippe Vernon, Johan J. van Tol, Yunga Wu, Donghui Wu, Zhijing Xie, Bettina Weber, Paul Illmer, Julia Seeber

## Abstract

Despite the importance of healthy soils for human livelihood, wellbeing, and safety, current gaps in our knowledge and understanding of biodiversity in soil are numerous, undermining conservation efforts. These gaps are particularly wide in mountain regions where healthy soils are especially important for human safety and yet evidence is accumulating of ongoing degradation, posing significant threats to ecosystem functioning and human settlements.

To analyse these gaps in detail, we synthesise current research on the global diversity of microorganisms, cryptogams, and invertebrates in mountain soils above the treeline. This synthesis is based on a semi-quantitative survey of the literature and an expert-based analysis. Our work reveals not only deficiencies in geographic cover but also significant gaps in taxonomic coverage, particularly among soil protists and invertebrates, and a lack of (functional and ecological) description of the uncultivated majority of prokaryotes, fungi, and protists. We subsequently build on this overview to highlight opportunities for research on mountain soils as systems of co-occurring species that interact in complex environmental matrices to fulfil critical functions and make essential contributions to life on land.

Closing gaps in biodiversity research in mountain soil is crucial to enhance our understanding and to promote laws and guidelines advancing international soil biodiversity conservation targets in mountains. Addressing sparse and biased data, recognizing the impact of environmental changes on mountain ecosystems, and advocating dedicated policies are essential strategies to safeguard mountain soils and their biodiversity.

**GLOSSARY:** 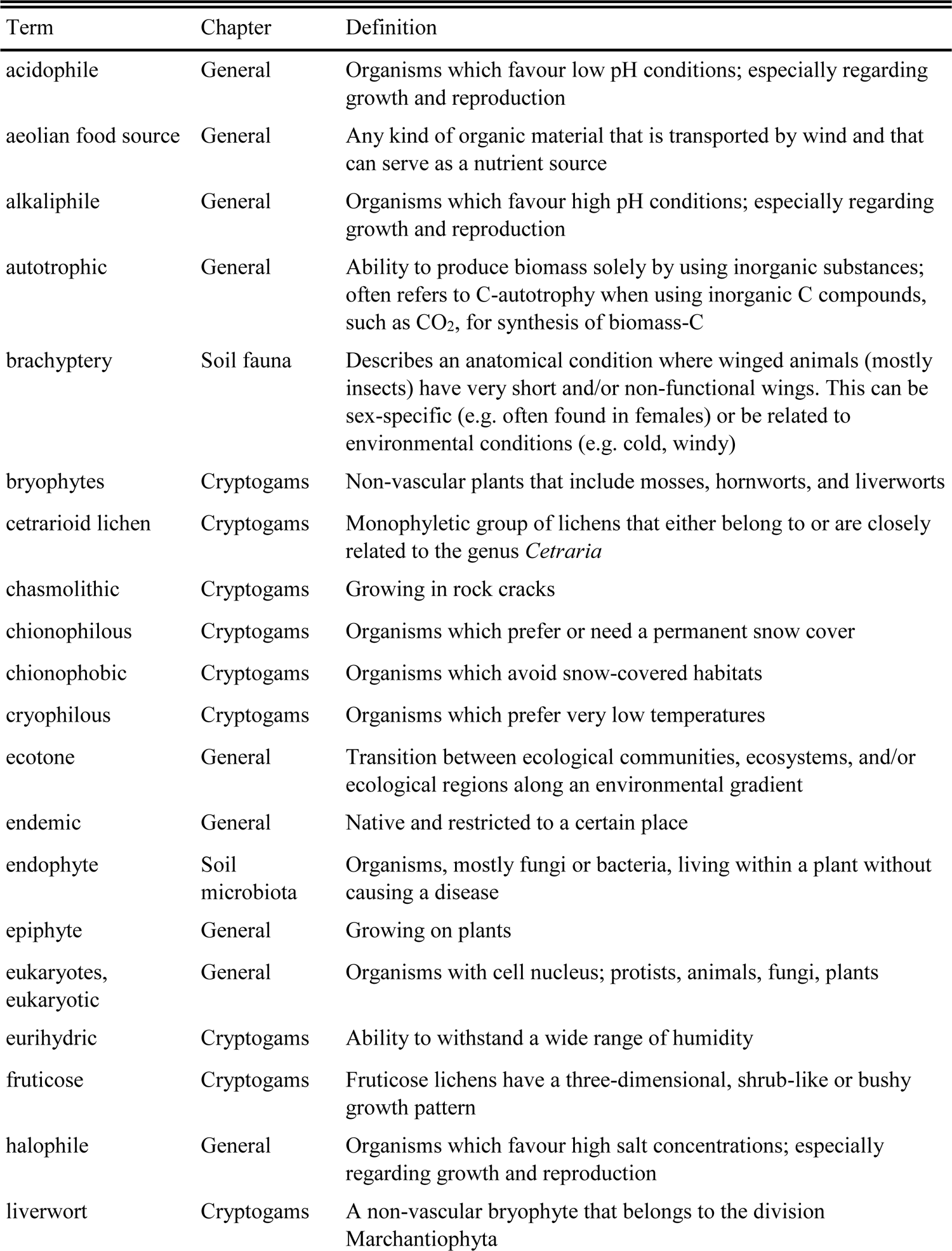

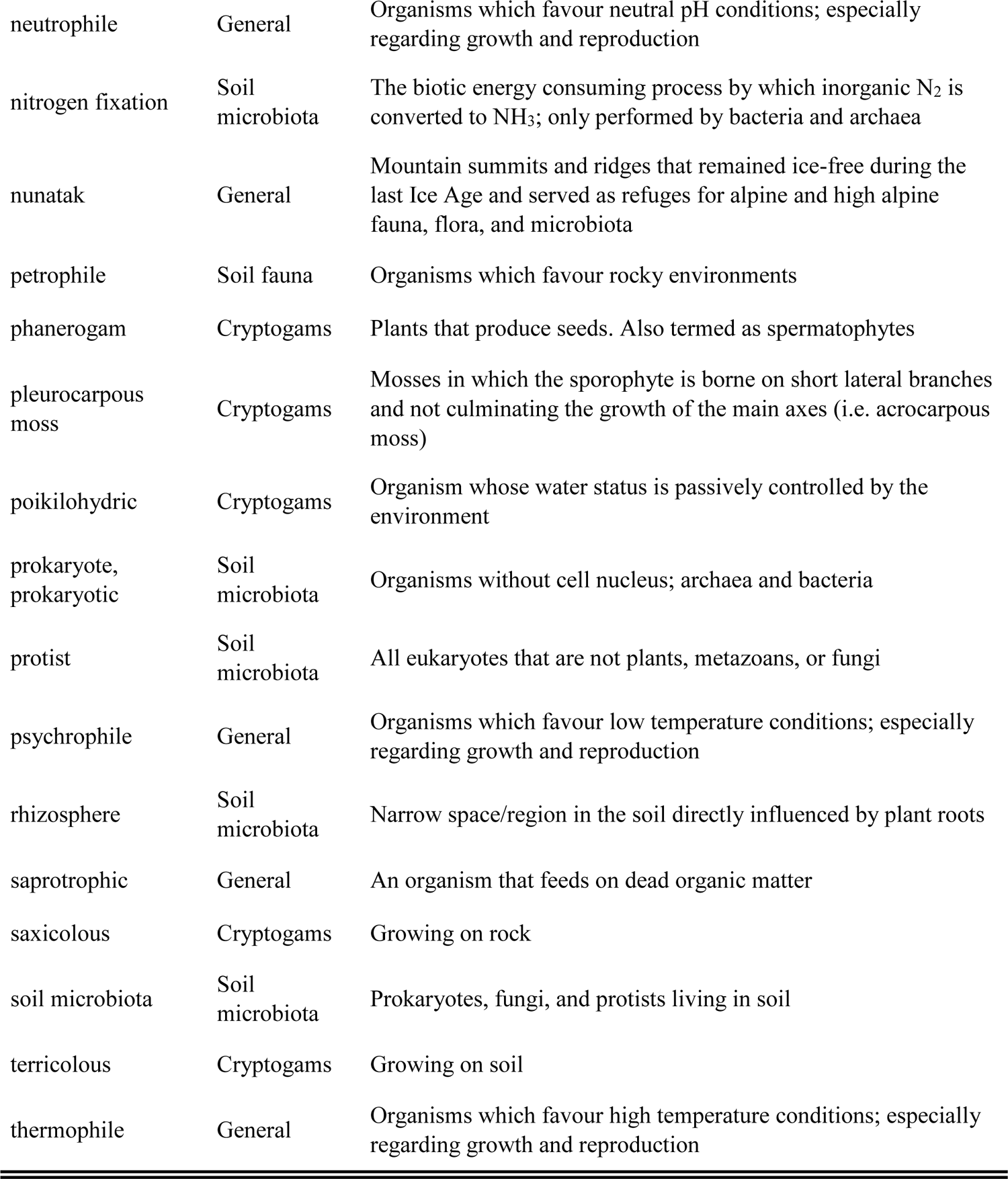

## I. INTRODUCTION

In recent years, our awareness of the importance of soils and their biodiversity has steadily increased, pressed by the growing evidence of rapid soil degradation worldwide and across all biomes (European Environment Agency, 2019; FAO *et al*., 2020; Anthony, Bender & van der Heijden, 2023). Because of their environmental, societal, and economic consequences, soil degradation and the loss of soil biodiversity pose a major threat to humankind. The need to protect soils has become an international priority (IPBES, 2018), reflected in both the Agenda for Sustainable Development of the United Nations and the recently adopted Kunming-Montreal Global Biodiversity Framework (United Nations, 2015; UN Convention on Biological Diversity, 2022). The demand for data, knowledge, and global responses to the challenge of how to safeguard soils and their biota has led to an increasing number of international initiatives, including the Soil Biodiversity Observation Network (SoilBON), the Global Soil Biodiversity Initiative (GSBI), the International Network on Soil Biodiversity (NETSOB), and the Global Soil Laboratory Network (GLOSOLAN). All these initiatives aim at providing the biological and ecosystem information needed to implement sustainable management and conservation of soils.

Despite these recent developments, major gaps and blind spots still exist in soil research and in available data and knowledge on soil biodiversity. This is particularly the case with mountains (Baruck *et al*., 2016; Guerra *et al*., 2020), even though mountain soils are critical for many ecosystem processes, functions, and services, and their maintenance and stability are particularly important in terms of hazards and natural risk management (FAO, 2015; Stanchi *et al*., 2023). Given that mountain soils can take thousands of years to develop (up to 1000 years for 2–3 cm in (high) alpine areas (Stanchi *et al*., 2023), their degradation and gradual erosion as a result of overexploitation and poor management may ultimately lead to a loss of biodiversity and associated ecosystem collapse, with no option for recovery (Körner, 2021; Singh *et al*., 2023). It emphasises the complexity of ecological restoration, pointing out that while repairing functions and maintaining existing taxa is feasible to some extent, the irreversible loss of certain locally adapted species, especially in isolated environments like nunataks, is a significant concern. These threats are further exacerbated by climate and land-use change, as well as the increasing occurrence of invasive non-native species (Palomo, 2017; Zucconi & Buzzini, 2021; Iseli *et al*., 2023). Here, we first provide a synthetic overview of the current state of knowledge on biodiversity in mountain soils above the treeline. We subsequently build on this overview to highlight opportunities for research on mountain soils as systems of co-occurring species that interact in complex environmental matrices to fulfil critical functions and make essential contributions to life on land. We restrict this review to alpine soils above the treeline in mountain regions (Fig. 1, Table 1). The term ‘alpine’ in this context specifically refers to soils located in mountainous areas above the treeline. The synthesis was performed as a collaborative effort by members of the Global Mountain Biodiversity Assessment (GMBA) ‘Mountain Soil Biodiversity’ working group, who summarised current literature in their respective fields of expertise. This body of literature was further consolidated based on a semi-comprehensive review of available publications.

**Fig. 1.**
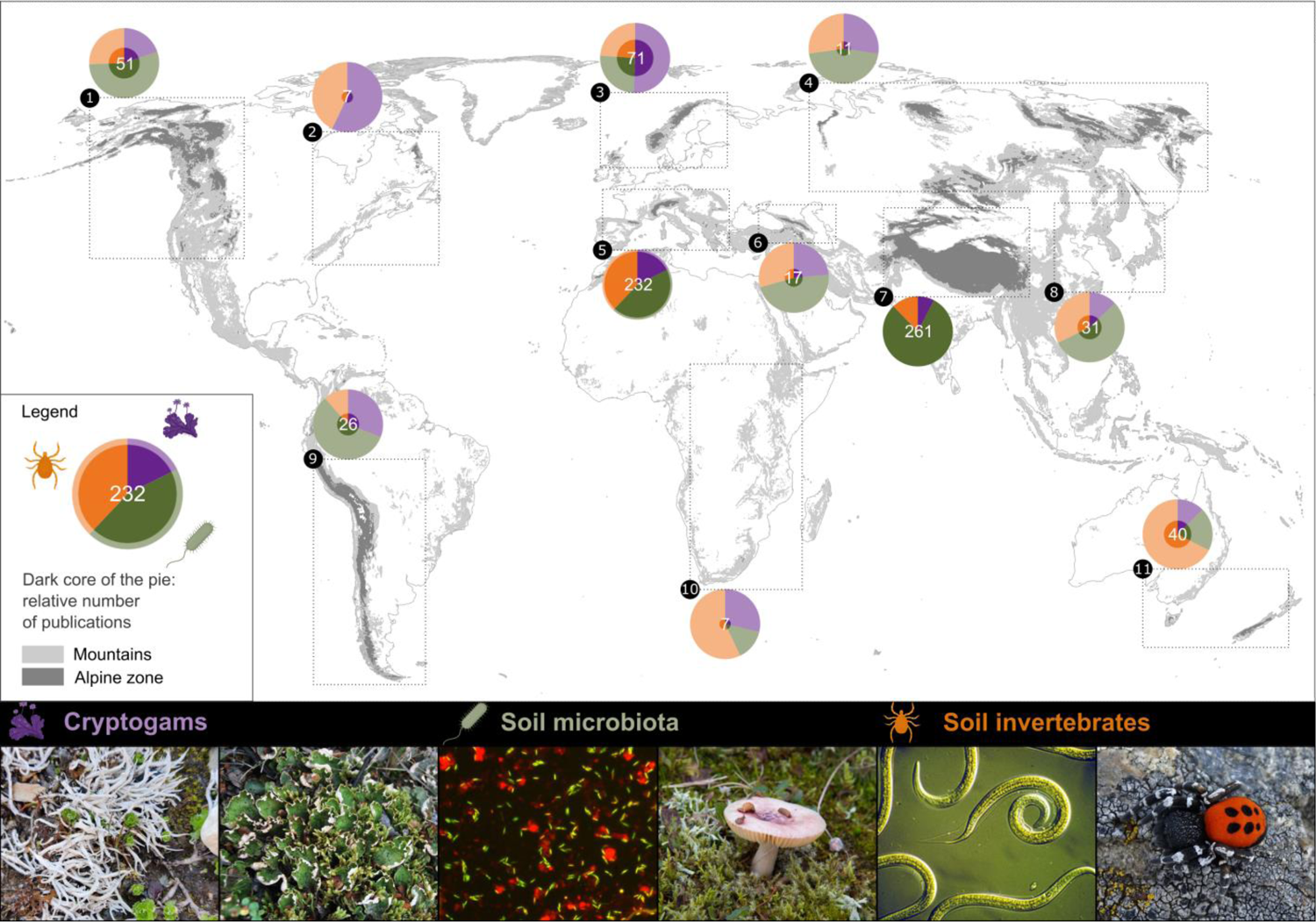
Global map of the number of scientific papers on biodiversity in mountain soils above the treeline (cryptogams, soil microbiota, and soil invertebrates) by alpine region. The dark core of the pies represents the number of publications in the respective area compared to the number of the region with the most publications (i.e. Central Asia). The alpine regions were named here as (1) North American Cordillera, (2) Appalachians & Northeast Ranges, (3) Northern Europe, (4) North Asia, (5) Central & Southern Europe, (6) Caucasus, (7) Central Asia, (8) East Asia, (9) Andes & South America, (10) Afro-alpine Region, and (11) Australia & New Zealand. See Fig. S1 for an alternative version showing the density of papers per km² of alpine area. Photos from left to right: Cryptogams: Arctic-alpine lichen *Thamnolia vermicularis*, arctic-alpine lichen *Peltigera aphthosa* (credit: Bettina Weber); Soil microbiota: DNA stained microscope preparation of soil bacteria (credit: Nadine Praeg & Paul Illmer), *Russula* sp. (credit: Andrea J. Britton); Soil invertebrates: Nematodes and a male velvet spider *Eresus sandaliatus* (credits: CSIRO Entomology and Michael Steinwandter, respectively).

**Fig. 2.**
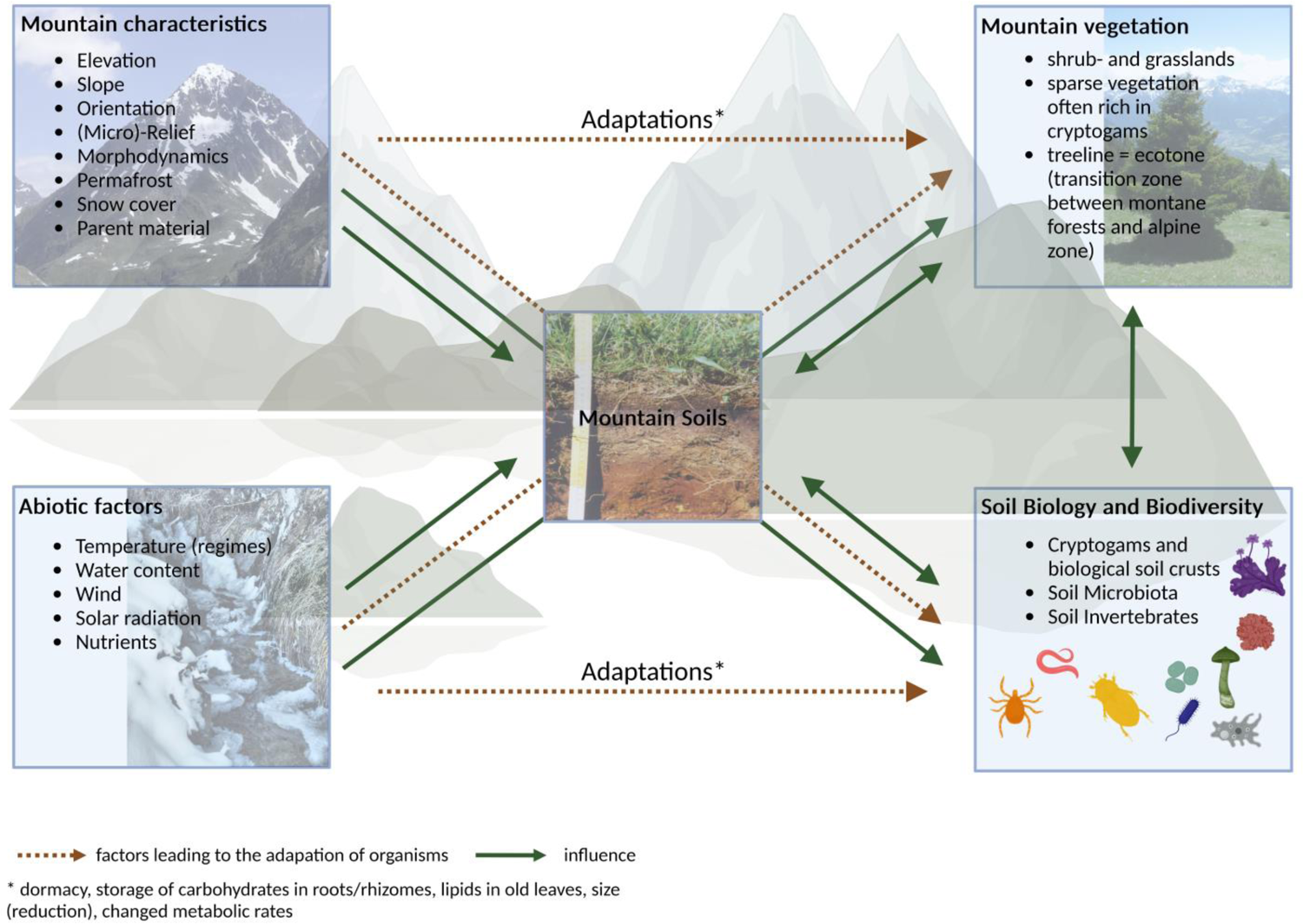
Overview of mountain characteristics and abiotic factors, influencing alpine landscape, mountain soils, and soil organisms.

**Table 1.** List of the alpine mountain regions included in this review, presenting their respective area, and the corresponding number of papers per main soil organism group. The identification (ID) numbers of the alpine mountain regions align with those referenced in the Figs. 1 & 3–5. The designation of the regions identified as ‘alpine region’ along with a summary of the corresponding mountain ranges is detailed in Table S3. For visual comparison of the areas (tree map) of the alpine mountain regions, please refer to Fig. S2 in Appendix S1.

| Nr. | Alpine region | Area [km²] | Number of scientific articles |  |  |  |
| --- | --- | --- | --- | --- | --- | --- |
|  |  |  | Cryptogams | Microbes | Invertebrates | Total |
| 1 | North American Cordillera | 306,815 | 10 | 28 | 13 | 51 |
| 2 | Appalachians & Northeast Ranges | 10,929 | 4 | 0 | 3 | 7 |
| 3 | Northern Europe | 47,623 | 36 | 18 | 17 | 71 |
| 4 | North Asia | 395,738 | 3 | 5 | 3 | 11 |
| 5 | Central & Southern Europe | 26,606 | 41 | 103 | 88 | 232 |
| 6 | Caucasus | 16,567 | 4 | 8 | 5 | 17 |
| 7 | Central Asia | 2,188,571 | 20 | 209 | 32 | 261 |
| 8 | East Asia | 2,778 | 4 | 17 | 10 | 31 |
| 9 | Andes & South America | 547,091 | 8 | 15 | 3 | 26 |
| 10 | Afro-alpine Region | 497 | 2 | 1 | 4 | 7 |
| 11 | Australia & New Zealand | 12,639 | 5 | 8 | 27 | 40 |
|  | Sums | 3,555,854 | 137 | 412 | 205 | 754 |

## I. METHODS

The synthesis was performed by querying ‘Web of Science’ for scientific papers on biodiversity in mountain soils in the alpine zones of the world (i.e. above the treeline in temperate and continental climatic zones, excluding tropical areas) and attributing each paper to one or more of the organismic groups considered here, making a distinction between primary and secondary focus as many publications covered more than one organismic group (Figs. 3–5) (see detailed methods in Appendix S1). Each paper was also attributed to one of the 11 defined alpine regions (see Table 1 and Table S3) according to the mountain ranges or systems that were specified in the title, abstract, keywords, or methods. The lists were subsequently reviewed and validated and supplemented with references from the authors personal reference databases. See Appendix S1 for a detailed description of the literature search and data processing.

**Fig. 3.**
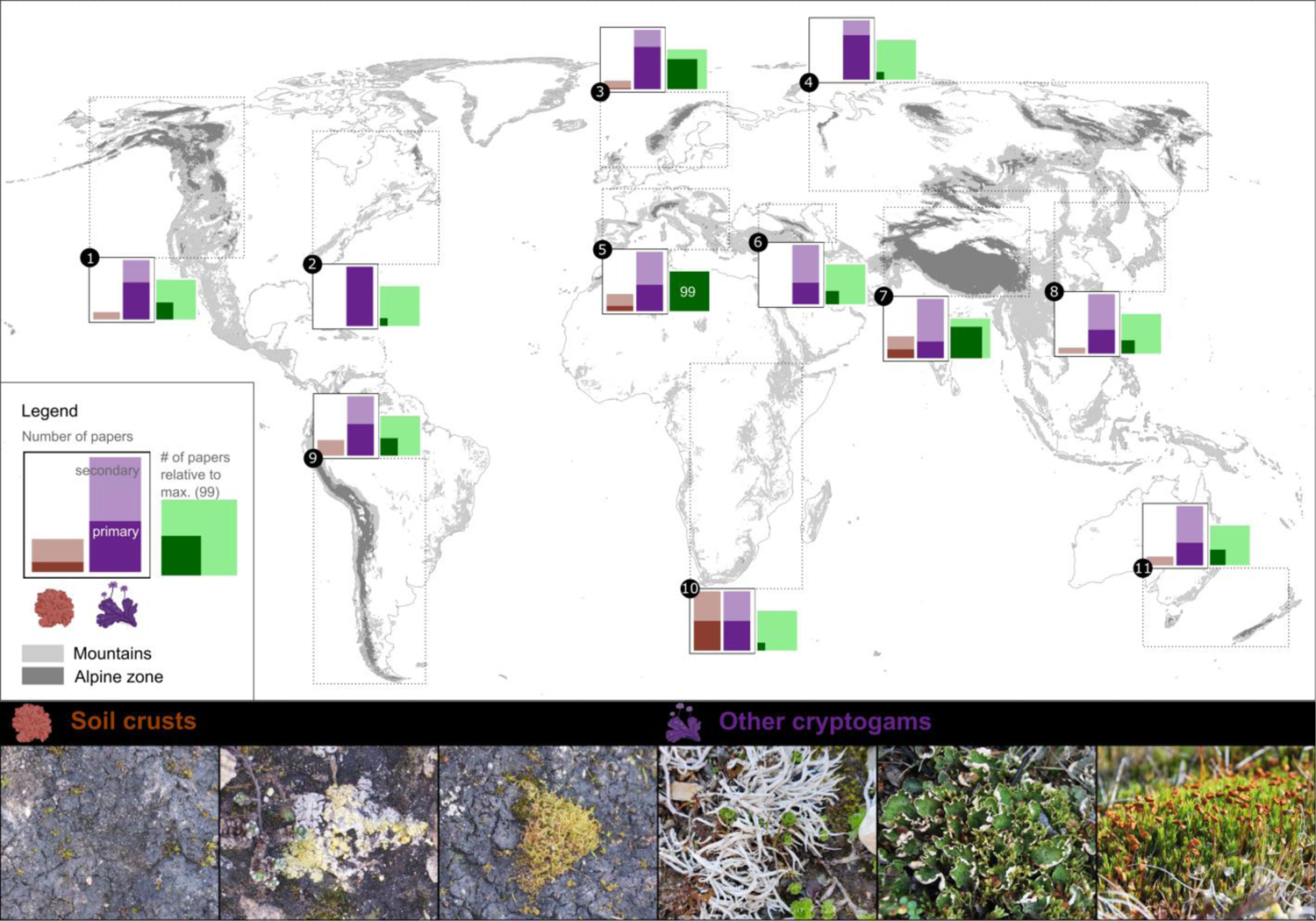
Global map of available scientific articles focusing on cryptogams in alpine mountain soils. Number of publications is given per crust type and relative to the maximum of papers found (Central & Southern Europe): the dark-coloured part of the bar represents those papers where the organismic group was likely the primary object, the light-coloured part represents those papers where the organismic group was mentioned, but not as the main subject of the publication. See Appendix S1 for a detailed description of the methods and full lists of publication numbers per region and soil organism group. Photos from left to right: Soil crusts: cyanobacteria-dominated soil crust (dark surface colouration) intermingled by bryophytes, in alpine zone of Großglockner, Austria (credit: Stefan Herdy); cyanobacteria-dominated biocrust mixed with chlorolichens, dominated by *Fulgensia* sp., alpine zone of the Großglockner, Austria (credit: Stefan Herdy); cyanobacteria-dominated biocrust mixed with mosses, dominated by *Tortella* sp., alpine zone of the Großglockner, Austria (credit: Stefan Herdy); Other cryptogams: arctic-alpine lichen *Thamnolia vermicularis,* alpine zone of the Großglockner, Austria (credit: Stefan Herdy); arctic-alpine lichen *Peltigera apthosa* in vicinity of Kangerlussuaq, Greenland (credit: Bettina Weber); moss *Polytrichum* sp. in vicinity of Kangerlussuaq, Greenland (credit: Bettina Weber).

**Fig. 4.**
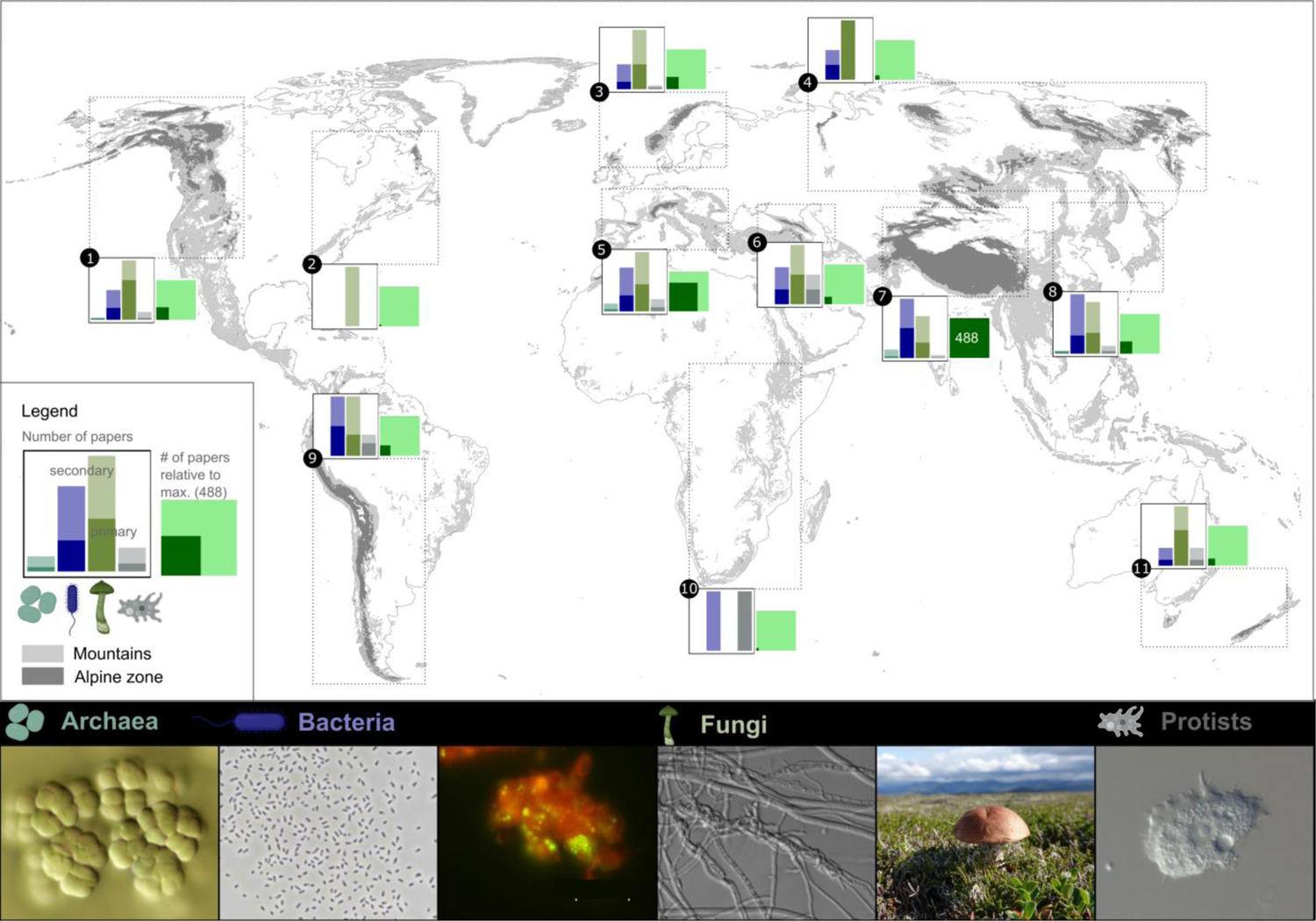
Global map of available scientific articles focusing on microbial diversity in alpine mountain soils. Number of publications are given per microbial group and relative to the maximum of papers found (Central Asia): the dark-coloured part of the bar represents those papers where the organismic group was likely the primary object, the light-coloured part represents those papers where the organismic group was mentioned, but not as the main subject of the publication. See Appendix S1 for a detailed description of the methods and full lists of publication numbers per region and soil organism group. Photos from left to right: Archaea: *Methanosarcina* sp. (credit: Paul Illmer), Bacteria: *Methylosinus sporium* (credit: Nadine Praeg), DNA (green) stained microscope preparation of soil bacteria attached to soil particle (red) (credit: Nadine Praeg & Paul Illmer), Fungi: *Trichoderma asperellum* intercoiled with *Botrytis* sp. (credit: Siebe Pierson), *Leccinum vulpinum* (credit: Andrea J. Britton), Protists: *Acanthamoeba* sp. (credit: Kenneth Dumack).

**Fig. 5.**
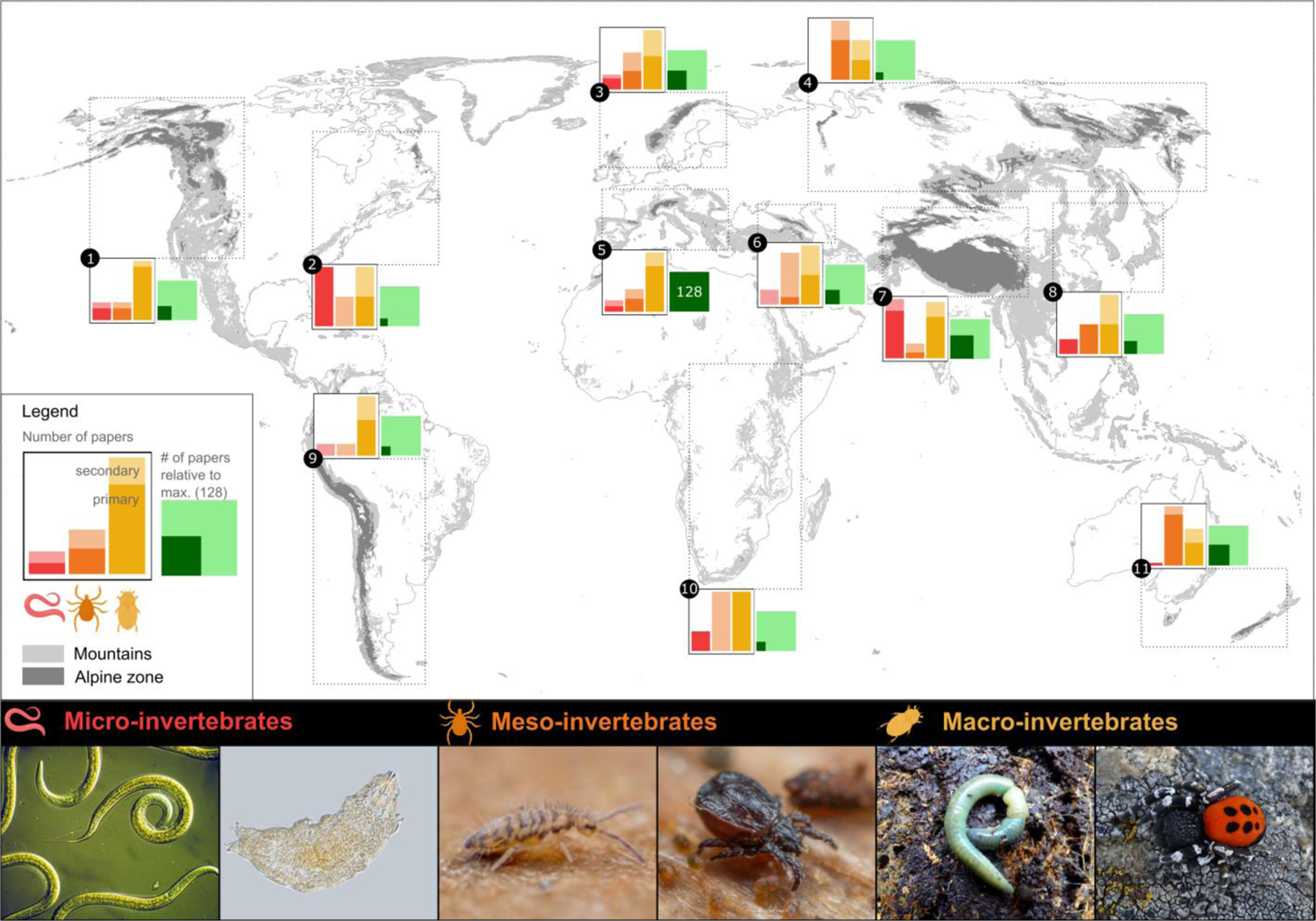
Global map of scientific articles focusing on invertebrates in alpine mountain soils. Number of publications are given per invertebrate group and relative to the maximum of papers found (Central & Southern Europe): the dark-coloured part of the bar represents those papers where the organismic group was likely the primary object, the light-coloured part represents those papers where the organismic group was mentioned, but not as the main subject of the publication. See Appendix S1 for a detailed description of the methods and full list of publication numbers per region and soil organism group. Photos from left to right: Micro-invertebrates: images of Nematoda (credit: CSIRO Entomology), and a Tardigrada *Macrobiotus* sp. (credit: Michala Tůmová) through microscopes; Meso-invertebrates: the Collembola *Entomobrya nivalis* and the Acari *Platynothrus pelifer* (credit: both Frank Ashwood); Macro-invertebrates: the ‘green’ earthworm *Aporecctodea smaragdina* inhabits calcareous mountain soils in the European Alps and Dinaric Alps, male velvet spider *Eresus sandaliatus* found in alpine dry pastures in the Central European Alps (credit: both Michael Steinwandter).

## II. CHARACTERISTICS OF MOUNTAIN ZONES ABOVE THE TREELINE

### (1) Soil

Major soil types occurring in the alpine and nival zones of temperate and continental mountains include (according to the soil taxonomy, Soil Survey Staff, 2022) Entisols, Inceptisols, Mollisols, Histosols, permafrost-affected soils, and Podzols, the latter mainly found on siliceous rocks, slopes with conifers, and alpine dwarf-shrub zones (Gelisols) (Price & Harden, 2013). Soil formation in mountain areas is – besides climatic factors – typically controlled by microrelief and morphodynamics, gravitational and fluvial dynamics, solifluction, and wind-related processes (Egli, Dahms & Norton, 2014). Accordingly, soil types and properties show small-scale heterogeneity (Egli & Poulenard, 2016) resulting from the high variability in (micro-) climate, topography, orientation, slope, and regional/local wind systems (Burga, Klötzli & Grabherr, 2004; Hoorn, Perrigo & Antonelli, 2018). Properties that are specific to alpine soils include an incomplete development (Donhauser & Frey, 2018), with slow humus accumulation and limited nutrient supply, even though accumulation of wind-blown fine material may, on a small scale, improve the physical and chemical characteristics of stony substrates (Gild, Geitner & Sanders, 2018). Detailed information on mountain soil types and characteristics is an important prerequisite to understanding biodiversity in soil and the unique adaptations of soil (micro-)organisms to overall hostile environmental conditions (Fig. 2) (Pellissier *et al*., 2014; Orgiazzi *et al*., 2016; Yashiro *et al*., 2016; Mod *et al*., 2020; Seppey *et al*., 2020). However, this information remains rare (Baruck *et al*., 2016; Guisan *et al*., 2019) and needs to be implemented through specific and targeted initiatives (e.g. the establishment of the Alpine Soil Partnership in 2017).

Generally, life in mountain soils is determined by their abiotic properties, including water content and temperature as the main drivers of chemical and physical weathering, parent material, including chemical composition, physical properties, resistance against weathering, and the predetermination of soil pH as well as organic matter quality and quantity (Fig. 2) (Paul & Clark, 1996). Both the length of the growing season and the mean temperature in mountain soils typically decrease with increasing elevation. In eastern Scotland, for example, Britton *et al*. (2011) measured a decrease in mean summer soil temperature at 10 cm depth from 9.5 °C at the treeline at 640 m a.s.l. to 7.2 °C at 908 m and a decline in the length of the growing season from 198 to 156 days based on a threshold of 5 °C. The low temperatures typical of mountain soils at high elevations favour the accumulation of organic matter through reduced decomposition rates, which can result in large amounts of soil carbon (Praeg *et al*., 2020). Accordingly, mountain regions have received increasing attention for their contribution to terrestrial C storage (Hagedorn, Gavazov & Alexander, 2019; Stanchi *et al*., 2021). While nival (and alpine) soils exhibit minimal C stocks (Frey *et al*., 2016; Adamczyk *et al*., 2019; Luláková *et al*., 2019), soils at lower elevations, where temperatures and plant coverage are higher, may act as larger C sinks.

An additional factor affecting life in mountain soils is the presence of a substantial and long-lasting snow cover. Together, seasonal snowpack depth, duration, and melt-out control the onset and duration of the growing season, mitigate low soil temperatures, and affect microbial activity, soil nutrient cycling, soil gas fluxes, and pedogenesis (Freppaz *et al*., 2017). These factors further determine community composition, and their high spatial variability may enable the close co-occurrence of species adapted to differing environmental conditions such as chionophilous and chionophobic taxa (Odland & Munkejord, 2008; Carlson *et al*., 2015; Niittynen & Luoto, 2018; Seeber *et al*., 2021; Panchard *et al*., 2023). In addition to the presence of substantial (and natural) snow cover, further drivers of change in abiotic soil properties include winter sports operations. These activities, such as the establishment and maintenance of (large) ski areas, including levelling and grading operations, represent strong mechanical disturbance. This promotes the breakdown of soil aggregates, causes the exposure of organic matter that was previously protected in undisturbed soils (Gros *et al*., 2004), and fosters soil erosion, which all together lowers the organic C content (Delgado *et al*., 2007; Negro *et al*., 2013) and reduces the soil micropore porosity, with consequences for soil aeration and water holding capacity (Pohl *et al*., 2009). Additionally, artificial snowmaking uses nucleating agents and water, often diverted from lakes and streams, which contain mineral and organic compounds that are not present in natural snow. This provides an additional input of solutes during snow melting (Wipf *et al*., 2005; Roux-Fouillet, Wipf & Rixen, 2011), resulting in higher soil pH and electrical conductivity (Delgado *et al*., 2007; Freppaz *et al*., 2013; Casagrande Bacchiocchi *et al*., 2019).

Considering the impact of global change, mountain soils are likely to undergo major transitions. Nival soils, as stated above, can be expected to increasingly serve as C sinks with climate warming. They are equally expected to become N sinks due to increased plant productivity under warmer conditions (Steinbauer *et al*., 2018). This holds especially true for barren or sparsely vegetated soils with low C and N content, where increased plant growth and primary production clearly outweigh temperature-driven increases in soil respiration (Hagedorn *et al*., 2019). In contrast, distinct increases in C loss have also been reported as a consequence of increased temperatures allowing lowland plants to colonise alpine environments (Walker *et al*., 2022). Thus, predictions about the amount and the dynamics of C and N cycling in response to global warming can only be made on a basis of a better understanding of the complex biotic and abiotic interactions in mountain soils (Fig. 2).

### (2) Vegetation

Mountain ecosystems above the treeline consist of alpine shrub- and grasslands that gradually give way to high alpine areas extending into the zone of perennial snow and ice. These alpine areas typically have sparse vegetation but often rich cryptogam communities. The treeline itself is a transition zone, a so-called ecotone, between the higher montane forest, often dominated by coniferous trees, and the alpine zone.

The abundance of plant species and their distribution are determined by temperature, water availability, and the duration of snow cover, which results from the interacting effects of temperature, precipitation, topography, and wind (Rodwell, 1992a, 1992b; Thompson & Brown, 1992; Panchard *et al*., 2023). Alpine grasslands share many structural and functional traits and characteristics with polar grass-dominated tundra ecosystems (Riebesell, 1982; Janišová *et al*., 2011; Dengler *et al*., 2014), and in both systems, low air- and soil temperatures are an important factor for plant growth. Frost often limits the growth of trees and shrubs (Peel, Finlayson & McMahon, 2007), whereas permafrost (i.e. continuous frost conditions) controls the entire soil system and slows down all biotic activity (Parolo & Rossi, 2008; Zollinger *et al*., 2013; Goordial *et al*., 2016; Giaccone *et al*., 2019; Jin *et al*., 2020). At the upper limit of grasslands, occasionally increased aridity and shortened vegetation periods cause poikilohydric cryptogamic organisms to gradually replace the standing euryhydric phanerogams (Körner, 2021).

Adaptations of alpine vegetation to short and cold growing seasons include the ability to metabolise rapidly at low temperatures, the transition to dormancy as a strategy to withstand the rigours of winter, and the storage of carbohydrates in roots/rhizomes or of lipids in old leaves for regrowth and flower primordia formation (Billings, 1974) (Fig. 2). Alpine plants are also adapted to intense solar radiation, as well as to extended periods of dehydration. Whereas the structure and composition of alpine vegetation depends on soil type and the chemical and physical properties of soil, plant communities, in turn, influence soil structure, properties, and stability.

## III. CRYPTOGAMS AND BIOLOGICAL SOIL CRUSTS

#### KEY ASPECTS

- Cryptogam communities are widely distributed across different elevational zones in alpine regions
- Biological soil crusts are mainly restricted to high alpine areas.
- Lichen and bryophyte diversity and productivity decline above the treeline towards the nival belt, but at slower rates than that of vascular vegetation.
- The composition of lichen and bryophyte communities is strongly related to bedrock chemistry and soil texture.
- Climate change causes bryophytes to move upwards, whereas at lower elevations sensitive and rare lichens and bryophytes are endangered by vascular plant growth.
- We found 137 publications dealing primarily with alpine cryptogams (i.e. 16 for biological soil crusts and 121 for cryptogams in general), mainly from the mountain regions of Central & Southern Europe (29.9%), Northern Europe (26.3%), and Central Asia (14.6%); see Fig. 1, 3, and Table S4.

### (1) Cryptogams

Cryptogams are non-vascular organisms that do not form flowers and seeds but reproduce by fission, fragmentation, and spores. They comprise lichens, bryophytes, eukaryotic algae, and cyanobacteria (Büdel, Friedl & Beyschlag, 2023). Cryptogams occur widely at different elevations in alpine regions, where they grow epiphytically on vascular plants as well as on and within rocks (saxicolous) and on soil. In some cases, soil-inhabiting (epigaeic) organisms form biological soil crusts (abbreviated as biocrusts). Since biocrusts are defined as an ‘intimate association between soil particles […] and organisms which live within, or immediately on top of, the uppermost millimetres of the soil’ (Weber *et al*., 2022), they do not include fruticose lichen and bryophyte carpets, which mainly grow above the soil and form valuable vegetation components on their own. A detailed description of alpine biocrusts is given in Chapter IV.2 below.

In Europe, descriptions exist for 200 lichen species in the nival belt of the Alps, with a remarkable development of the genera *Cetraria*, *Parmelia*, and *Umbilicaria* (Ozenda & Borel, 2003). The number of bryophytes and macrolichens increases towards the North, with 439 species of bryophytes in the Italian Alps (Pedrotti & Grafta, 2003), 65 bryophyte and 218 lichen species in the south-eastern Carpathian Mountains (Coldea, 2003), 150–200 species of lichens in the Pyrenees (Gómez, Sesé & Villar, 2003), and about 558 bryophyte species in the Southern and Northern Scandes (Virtanen *et al*., 2003). Areas with the highest species richness of bryophytes but also the highest numbers of threatened species, are located in the eastern European Alps, Carpathian Mountains, eastern Pyrenees, and the Scandes (Hodgetts *et al*., 2019). The highest elevational records for lichens and bryophytes are found in the subtropical Dry Andes, near the summit of Socompa volcano at 6,060 m (Halloy, 1991) and on Mount Everest, Himalaya at 7,400 m (*Lecidea vorticosa* (FLÖRKE) KÖRB. and *Pertusaria bryontha* (ACH.) NYL.; (Miehe, 1988).

Grassland ecosystems contain a high diversity of cryptogamic autotrophs, especially bryophytes and lichens, whose occurrence is primarily determined by elevation and exposition (Cleavitt, 2004; Daniëls *et al*., 2004; Baniya *et al*., 2012; Rai, Upreti & Gupta, 2012). Their diversity and productivity follow the same elevational gradients of temperature and aridity as phanerogams (Sundstøl & Odland, 2017); species richness of lichens, bryophytes, and algae increase above the treeline and progressively decline towards the nival belt (Austrheim, 2002; Vittoz *et al*., 2010).

In temperate central European mountains, alpine grasslands are characterised by the presence of large meadows dominated by genera including *Festuca* and *Carex* (Ozenda, 1988), driven by the presence of a long-standing, deep snow cover, which melts relatively fast in spring. In this region, communities dominated by bryophytes and lichens cover only relatively small areas. In contrast, in Central Asian and Scandinavian mountains, higher aridity or a stronger clearing of snow cover by wind promotes communities richer in cryptogams and less dominated by graminoids. This trend is also observed in continental Nearctic mountain ranges such as the Rocky Mountains (Leuschner and Ellenberg 2017). In all mountain ranges, snow abrasion poses a major mechanical challenge to alpine vegetation of exposed habitats (Wieser, Holtmeier & Smith, 2014) and promotes the development of stress tolerant cryptogamic communities in windswept localities.

The topographic heterogeneity found in high alpine slopes tends to intersperse grasslands with azonal communities of saxicolous, terricolous, and chasmophytic cryptogams that colonise skeletal soils and patches of exposed mineral substrate. These azonal patches become more abundant towards the nival zone and in arid or windswept localities. There, some fruticose lichens and pleurocarpous mosses grow among graminoid patches and are thereby an integral part of alpine grasslands. The lower dependence of cryptogams on substrate presence can create highly discordant diversity patterns between cryptogams and vascular plants, as well as among cryptogam groups (Di Nuzzo *et al*., 2021). However, moss and phanerogam richness also showed a strong correlation with soil richness and diversity on the Hardangervidda plateau in Norway (Vestvidda, Southern Scandes), whereas there was no correlation with liverworts (Odland, Reinhardt & Pedersen, 2015). In Palearctic and Nearctic mountains, *Cladonia* species and pleurocarpous mosses such as *Pleurozium* sp. tend to dominate towards the treeline, while cetrarioid species or *Thamnolia* sp. become more common towards the upper part of the gradient.

The exact composition of lichen communities varies significantly, depending on the bedrock chemistry and resulting texture of the mineral fraction of local soils (Guo & Cao, 2001). Calcium-rich spring seeps were observed to form a refugium harbouring a rich variety of bryophytes and lichens (Miller, Fryday & Hinds, 2005). Conversely, areas with higher water retention and permanently flooded soils develop into bryophyte dominated bogs mostly shaped by *Sphagnum* species and pleurocarpous mosses (Halsey, Vitt & Gignac, 2000; Wahren, Williams & Papst, 2001; Bragazza, Gerdol & Rydin, 2003). Also in wetlands, fens, springs, and snowfields, cryptogams can reach high diversity (Dierssen & Dierssen, 2005; Cooper *et al*., 2010; Austin & Cooper, 2016). In alpine fellfields, cryptogams are a prominent component, with lichens covering up to about 50% of the surface area on the Beartooth Plateau, Montana and Wyoming (Greater Yellowstone Rockies; Eversman, 1995). Generally, lichen communities develop on substrates with little mechanical perturbation under changing hydration conditions. Wet and soft soils of glacier forefields are not colonised by lichens. Soils in many windswept localities are typically colonised by fruticose lichens, which usually interlock with shrubby plants rather than being connected with the soil. In contrast to a common notion of lichens as pioneer vegetation, crustose lichens in biocrust communities need not only soil stability but also long stable time intervals to develop, and they can suffer extinction due to shading from nearby plants. One means by which evolution has made it possible for lichens to overcome competition with plants or unsuitable soil conditions is by the development of shrublike fruticose morphologies, which grow as lichen heath and pleurocarpous mosses between persistent plant vegetation such as dwarf shrubs (e.g. genera *Vaccinium, Salix,* and *Erica* with *Hypnum cupressiforme* HEDW.; Schellenberg & Bergmeier, 2020).

Compared to phanerogams, which show high levels of endemism, lichens associated with alpine grasslands have very broad distributions, often being apparently sub-cosmopolitan. This interregional connectivity in arctic-alpine organisms has been studied in lichens (Fernández-Mendoza & Printzen, 2013; Garrido-Benavent & Pérez-Ortega, 2017; Onuţ-Brännström, Tibell & Johannesson, 2017), but also occurs in bryophytes (Mirek & Piekos-Mirkowa, 1992), and seems to reflect range expansions originating from interregional connectivity during the Pleistocene. This does not exclude the rare endemism of lichens at higher elevations, which can be due to substrate conditions that are not or hardly found elsewhere, or to habitat shrinkage due to climatic factors (e.g. *Cetradonia linearis* (A.EVANS) J.C.WEI & AHTI is only found in few localities in the Appalachian Mountains; Woodward, 2021).

While soil properties are key in determining cryptogam presence and composition, effects are reciprocal, and cryptogams also influence soil properties. For example, in an alpine *Vaccinium* thicket accompanied by *Polytrichum strictum* MENZIES EX BRIDEL and *Sphagnum* sp., the moss cover caused a pedogenic feedback by increased water storage, which promoted stronger weathering and increased dissolved organic C contents in the soil. The latter then caused the soil to cross the threshold of podsolisation (Musielok *et al*., 2021). Similarly, mat-forming lichens have also been shown to influence litter decomposition and buffer soil temperatures in subalpine and alpine environments (van Zuijlen *et al*., 2020; Mallen-Cooper, Graae & Cornwell, 2021). Mosses, in turn, have been reported to mediate soil properties such as temperature, moisture, and C:N ratios, with varying effects depending on the shrub species under which they occur (Bueno *et al*., 2016). Generally, lichens and mosses also contribute to soil stability and reduce erosion in alpine environments (Martin *et al*., 2010).

The interactions between plants and lichens have also been explored (Favero-Longo & Piervittori, 2010). Most notably, the elimination of fruticose lichens, especially cetrarioid species, has been shown to significantly reduce the growth of neighbouring grasses and sedges (Jespersen, 2013), probably as a result of changes in microclimate, surface water retention, and protection from run-off. In an experimental approach, presence of most lichens facilitated seedling recruitment, while only very thick mats of *Cladonia stellaris* (OPIZ) POUZAR & VĔZDA had an inhibitory effect (Nystuen *et al*., 2019). Interactions between vascular plants and bryophytes are also variable. *Ptilium crista-castrensis* DE NOTARIS, a feather moss, was observed to have a negative effect on the survival of alpine tree seedlings, likely due to altered competition or nutrient availability, whereas *Sphagnum* mosses had no effect (Lett *et al*., 2020). In contrast, a study in a boreal forest-tundra ecotone (Central Labrador Ranges, Canada) revealed that a *Pleurozium schreberi* MITTEN seedbed improved seed emergence, survival, and nutrient availability for black spruce (Wheeler, Hermanutz & Marino, 2011). Similarly, *Racomitrium lanuginosum* BRIDEL, mats stimulated growth of the sedge *Carex bigelowii* TORR. EX SCHWEIN. in the alpine/subarctic tundra of Swedish Lapland (Carlsson & Callaghan, 1991).

Effects of environmental changes on cryptogams are likely diverse. Climate change in alpine regions generally causes a shift of bryophytes to higher elevations (Wen *et al*., 2022), whereas nutrient input, CO2 increase, or warming, cause vascular plant productivity to increase at the cost of sensitive and long-established soil lichen and bryophyte communities (Graglia *et al*., 2001; Klanderud, 2008; Dawes *et al*., 2017). Such changes in plant communities also affect microbial community composition, which may result in altered biogeochemical cycling (Bueno de Mesquita *et al*., 2017).

Anthropogenic drivers, such as grazing, trampling, and N deposition also affect cryptogams in complex and multiple ways. For instance, the effects of grazing are variable. Grazing in extensively farmed secondary grasslands has been shown to increase the diversity and coverage of bryophytes and lichens due to a decreased competition for light (Nascimbene, Fontana & Spitale, 2014). On the other hand, in mid-elevation pastures in India (3000–3400 m), where significant grazing by cattle occurs, lichen diversity is reduced compared to higher (3400–4000 m) and lower (2700–3000 m) habitats (Rai *et al*., 2012). In the Uinta Mountains of Utah (Western Rocky Mountains), grazing favoured the growth of crustose or squamulose lichens, whereas in ungrazed areas fruticose and foliose taxa also occurred (St. Clair *et al*., 2007). Vascular plants are also influenced, since further effects of (heavy) grazing include an increase in root biomass (Mayel, Jarrah & Kuka, 2021), also in alpine meadows (Yang *et al*., 2018), likely resulting from increased rates of nutrient cycling due to herbivore excretion. The effects of human trampling include the reduction of lichen abundance and diversity in an alpine heath ecosystem in northern Sweden (Jägerbrand & Alatalo, 2015) as well as a reduced coverage of the moss *Pleurozium schreberi* (WILLD. EX BRID.) MITT. in a subarctic grassland (Sørensen *et al*., 2009).

Finally, existing evidence for the effects of N deposition on cryptogams includes a loss in moss cover in an alpine *Racomitrium* moss-sedge heath in the United Kingdom (Britton *et al*., 2018), a general decline in richness, and a community shift from bryophytes and lichens towards graminoids (Nilsson *et al*., 2002; Armitage *et al*., 2014; Britton *et al*., 2019). Declines in moss cover are possibly due to the positive effects of N deposition on the growth of moss-associated fungi (Taylor *et al*., 2022). In a study in Norway, N addition caused a decrease in lichen cover and size (Fremstad, Paal & Mols, 2005), whereas in northern Sweden, N, P, and K fertilisation positively affected bryophyte biomass (Haugwitz & Michelsen, 2011).

After disturbance, succession under natural conditions or facilitated by restoration measures may help to reach the natural vegetation state again. In a study investigating the succession on alpine soil heaps in western Norway, it took about 30 years until the bryophyte and lichen cover and species richness was similar to the surrounding area (Rydgren *et al*., 2011). Similar results were also obtained in a separate study, where gamma diversity of cryptogams peaked 23 to 28 years after cessation of ploughing and fertilising subalpine grasslands (Austrheim & Olsson, 1999). In a study on habitat restoration after clearcutting of non-indigenous *Pinus mugo* TURRA in the Eastern Sudetes (Bohemian Massif), bryophyte diversity was mapped and compared to that in areas of undisturbed dwarf pine canopy and in autochthonous grassland areas. The results revealed a habitat homogenisation, as related to bryophytes, nine years after the impact, and suggested that restoration measures, in addition to clear-cutting, might be helpful to enhance restoration speed and quality (Zeidler *et al*., 2022). In a different study in Iceland, application of shredded turf led to a quick increase in bryophyte cover and thus might form a valuable restoration measure (Aradottir, 2012).

### (2) Biocrusts

As a pioneer community in alpine environments, biological soil crusts (biocrusts) comprise a dense layer of cyanobacteria, green algae, lichens, and bryophytes that covers the soil surface (Gold, Glew & Dickson, 2001; Huber *et al*., 2007; Karsten & Holzinger, 2014; Mikhailyuk *et al*., 2015; Weber, Büdel & Belnap, 2016) and grows in patches between vascular plants (Türk & Gärtner, 2003). Unlike lichen and bryophyte carpets, biocrusts do not elevate much above the soil surface. However, their presence and activity play a crucial role in forming soil aggregates, thereby enhancing soil stability. Early successional communities are dominated by cyanobacteria, which facilitate the gradual colonisation by lichens and bryophytes under suitable conditions. A more detailed definition of biocrusts and their delimitation against other cryptogam communities was published by Weber *et al*. (2022). Whereas cryptogams occur widely in alpine grasslands, biocrusts are mainly restricted to the high alpine zone, where they can achieve very large cover values. In the Austrian Alps (Hochtor, High Tauern), for example, biocrust coverage reached up to 30% of the surface area in the studied homogeneous vegetation unit (Büdel *et al*., 2014), with a high prevalence of cyanobacteria, which has also been observed in Himalayan soils (Reháková, Chlumská & Doležal, 2011). An increase with elevation was also detected for cyanobacterial biomass in cyanobacteria-dominated biocrusts in the Zanskar Range (Himalaya; Janatková *et al*., 2013).

The occurrence and composition of biocrusts in alpine regions appear to be mainly influenced by habitat availability and precipitation (Büdel *et al*., 2009; Lütz, 2012; Jung *et al*., 2018; Xiao *et al*., 2020). Biocrust activity status, in turn, is regulated by morphological, physiological, and local microclimatic conditions (Longton, 1988; Tamm *et al*., 2018). Additional variables affecting biocrust occurrence include elevation, aspect, snowpack, dust input in alpine areas, as well as standing vegetation (Miller, 2009; Sun *et al*., 2013; Mejia *et al*., 2020; Peer *et al*., 2022). Across four mountain ranges at high elevation in Ladakh, India, cyanobacterial occurrence along an elevational gradient from 3,700–5,970 m was mainly determined by the studied mountain range, but also elevation and vegetation type were relevant (Reháková *et al*., 2011). Whereas *Oscillatoriales* mostly occurred on alpine meadows, *Nostocales* were dominant in the subnival zone and screes.

Thawing of permafrost and glaciers produce particularly suitable habitats for biocrusts. Accordingly, apart from long-established cryptogam communities, (cyano-)bacteria, (lichenised) fungi, and algae play a key role in primary substrate colonisation after glacier retreat. This has been investigated in different alpine regions around the world, including Norway, Chile, Peru, and in the European Alps (Frey *et al*., 2013; Bilovitz *et al*., 2014b; Matthews & Vater, 2015; Krisai-Greilhuber *et al*., 2017). In Tierra del Fuego, Chile (Cordillera Darwin, Patagonian Andes), bacterial communities with cyanobacteria and algae of the order *Prasiolales* were the dominating groups close to the glacier terminus, whereas lichen-forming and parasitic fungi occurred in early successional stages (Fernández-Martínez *et al*., 2017). Cyanobacteria hosted by bryophytes and fertilising the immature soils by actively fixing atmospheric N were also observed 4–7 years after deglaciation (Arróniz-Crespo *et al*., 2014). Cyanobacteria were further described to play a vital role in primary succession with respect to both C and N fixation and soil stabilisation at high elevations (5,000 m) in the Cordillera de Vilcanota (Cordillera Oriental, Peru) (Schmidt *et al*., 2008). In the case of lichens, multiple studies in the European Alps describe an increasing abundance and diversity with moraine age (e.g. Bilovitz *et al*., 2014b, 2014a, 2015) and higher coverage compared to the surrounding non-glaciated area (Hestmark, Skogesal & Skullerud, 2007). In central Svalbard, repeated surveys of glacier forefields 10–20, 30–50, and 80–100 years after glaciation detected a marked shift in cryptogam community structure over time (Pessi *et al*., 2019).

Biocrusts provide ecosystem services via their functions in soil stabilisation, N and C fixation, nutrient accumulation, and water retention (Gold *et al*., 2001; Huber *et al*., 2007; Peer, 2010; Zheng *et al*., 2014; Jung *et al*., 2018; Borchhardt *et al*., 2019). They are further known for improving soil microenvironments, mainly due to the activity of microorganisms within the biocrust (Wei *et al*., 2022). In the case of glacier forefields, cyanobacteria fix and thus provide N to the strongly N-limited raw soils, with rates directly related to the availability of organic C (Wang *et al*., 2021). High N fixation rates by alpine *Collema*-dominated biocrusts in the mountains of Western Canada (i.e. Chilcotin Plateau (British Columbia Interior) and Southern Icefield Ranges (Saint Elias Mountains)) suggest an important contribution of cyanolichens to ecosystem N budgets (Marsh *et al*., 2006). While Antarctic and alpine biocrusts show similarities in composition, alpine biocrusts seem to be much more physiologically active than their polar counterparts (Colesie *et al*., 2014, 2016), with activity rates closely linked to the local climatic conditions (Raggio *et al*., 2017).

An additional effect of biocrusts is their influence on soil temperature. This was shown for instance on the Tibetan Plateau, where Ming *et al*. (2022) found that at a depth of 5 to 100 cm, soil temperatures were 0.6–1.0 °C lower in the presence of biocrusts; Xu *et al*. (2020) showed similar effects for moss-dominated biocrusts. These results differ from previous studies, where biocrusts increased surface temperatures due to their dark colour (Chamizo *et al*., 2013). A possible explanation is the high insulating potential of soil organic matter and the high water-holding capacity of the local biocrusts (Ming *et al*., 2022). Biocrusts on the Tibetan Plateau (i.e. Min Mountains and Qilian Mountains), were also observed to significantly reduce soil pH in the upper 10 cm (Xu *et al*., 2020) and to impact seed germination, thus influencing vascular plant community composition (Li *et al*., 2016). In another study, biocrusts tended to support the survival of *Nothofagus pumilio* (POEPP. & ENDL.) KRASSER tree seedlings in the southern Patagonian Andes (Pissolito, Garibotti & Villalba, 2021).

Besides their effects on environmental conditions, biocrusts also comprise various bacterial communities. Bacterial composition within biocrusts appears to be strongly impacted by the dominating photoautotrophs (i.e. cyanobacteria, algae, lichens, or bryophytes), whereas for microfungi such a link could not be observed (Maier *et al*., 2018; Abed *et al*., 2019). Along an aridity gradient on the Tibetan Plateau, algae-dominated biocrusts hosted more diverse bacterial communities, with diversity increasing with rising aridity, while in lichen-dominated biocrusts bacterial communities were less diverse and bacterial diversity decreased with rising aridity. Whereas the bacterial communities differed depending on the biocrust type, they were also influenced by environmental and stochastic processes (Wei *et al*., 2022). In alpine biocrusts, the soil–lichen interface was colonised by characteristic bacteria, namely Alphaproteobacteriota and Acidobacteriota (Muggia *et al*., 2013).

Adaptive strategies of lichens to severe conditions in alpine regions include accumulation of UV-absorbing phenolic usnic acid and storage of polyols for protection of cellular constituents during desiccation (Bligny & Aubert, 2012; Armstrong, 2017). Protective strategies of terrestrial, photosynthetic green algae include photoprotection, non-photochemical quenching and flexibility of secondary cell walls (Karsten & Holzinger, 2014; Kitzing, Pröschold & Karsten, 2014). Diurnal freeze–thaw cycles that frequently occur in high alpine habitats were shown to have no negative impact on the growth of cyanobacteria-dominated biocrusts collected in the Peruvian mountains of the Cordillera Oriental (Schmidt & Vimercati, 2019).

Land use by agriculture and recreational activities can cause severe damage to high-mountain ecosystems, including their biocrusts. On high elevational grasslands of the Ötztal Alps (European Alps) in Tyrol, Austria, for example, even weak trampling pressure caused a decrease in the frequency of sensitive species, including fruticose and crustose lichens (Grabherr, 1982). Also, in the Canadian Rockies of Alberta, recreational trails had substantially lower coverage of lichens and biocrusts, as compared to undisturbed sites (Crisfield, Macdonald & Gould, 2012). After disturbance of biocrusts in alpine habitats, restoration could be facilitated by inoculation with mature biocrusts (Letendre, Coxson & Stewart, 2019). Climate change poses another threat: In Switzerland, the observed increase in the mean elevation of bryophytes, driven by extinction of cryophilous species at lower elevations and by an upward movement at their upper range limits, was likely a result of recent climate change (Bergamini, Ungricht & Hofmann, 2009). Changes in bryophyte and lichen species richness, cover, and composition were also observed during a 15-year period from 2001 and 2015 in the Southern Scandes of Norway, with an effect on species interactions. For lichens, the observed decrease in species richness and cover over time was attributed to the increased competition with vascular plants (Vanneste *et al*., 2017).

## IV. SOIL MICROBIOTA

#### KEY ASPECTS

- Soil microorganisms are essential for mineralization processes, nutrient cycling and for many symbiotic relationships with plants and animals.
- The very great majority of soil microorganisms cannot be investigated by culture-based approaches that, therefore, need to be further improved.
- Knowledge about the enormous microbial diversity in alpine soils is distinctly increasing since the advent of high-throughput molecular methods.
- Microbial diversity is determined by complex interactions with abiotic soil properties such as soil pH, water content and quality and quantity of organic matter.
- Changes in microbial communities can have cascading effects on other components of the ecosystem.
- Fungal diversity is more strongly influenced by plants than the diversity of prokaryotes.
- Patterns of diversity and effects of abiotic and biotic drivers are distinctly group specific.
- We found 412 publications dealing primarily with alpine microbial soil diversity (i.e. 16 for archaea, 190 for bacteria, 184 for fungi, and 23 for protists), mainly from the mountain regions of Central Asia (50.7%), Central & Southern Europe (25.0%), and the North American Cordillera (6.8%); see Fig. 4 and Table S5.

### (1) Bacteria and Archaea (Prokaryotes)

Prokaryotes include two distinct phylogenetic domains, archaea and bacteria, which are both characterised by the absence of a cell nucleus. Most prokaryotes are unicellular and reproduce asexually. Due to a very high metabolic diversity (various chemo- and phototrophic ways of life), prokaryotes colonise almost every ecological niche on Earth.

A significant proportion of prokaryote diversity studies on high alpine soils (King *et al*., 2010; Yashiro *et al*., 2016) and alpine permafrost have been conducted on the Tibetan Plateau in Central Asia, which harbours the largest area of mountain permafrost soils globally (Cheng *et al*., 2022), resulting in 54 % of all prokaryote diversity studies in mountain soils being performed in Central Asia (Fig. 4). Studies addressing microbial diversity (all prokaryotes, selected studies also including fungi and protists) in mountain permafrost outside of China were conducted recently in the European Alps (Frey *et al*., 2016; Luláková *et al*., 2019; Praeg, Pauli & Illmer, 2019; Adamczyk, Rüthi & Frey, 2021; Sannino *et al*., 2021) and high-elevation soils from the Andes, Rocky Mountains, and Alaskan Brooks Range (Lipson & Schmidt, 2004; Nemergut *et al*., 2005; King *et al*., 2010; Ricketts *et al*., 2016; Wagner *et al*., 2017; Farrer *et al*., 2019). Despite partially harsh environmental conditions, mountain soils harbour a considerable bacterial diversity (Rime *et al*., 2015; Frey *et al*., 2016). Bacteria contribute substantially to biogeochemical cycles, both at the regional and supraregional scale (Donhauser & Frey, 2018) and together with archaea (and fungi, see Section V.2) are considered fundamental in stabilising soils and influencing the physical and biological development of soil ecosystems (Bernasconi *et al*., 2011). Prokaryotic colonisers contribute significantly to the initial build-up of biomass, by fixation of atmospheric CO2 and N2 (Frey et al., 2013), and using C and N from (microbial) necromass (Zumsteg, Schmutz & Frey, 2013b; Rime *et al*., 2016b; Donhauser *et al*., 2021). Quantification of precise amounts of CO2 and N2 fixation and usage of dead microbial cells as a non-negligible C pool in mountain soils is challenging due to the complexity and variability of mountain soils and is still lacking. Further nutrients, such as P and S, may be obtained from the bedrock by biological weathering (Frey *et al*., 2010; Brunner *et al*., 2011). As glaciers increasingly retreat with climate change, barren bedrock is exposed and colonised by pioneer microorganisms such as Acidobacteriota, Planctomycetota and Bacteroidota (Zumsteg *et al*., 2012; Rime *et al*., 2015; Rime, Hartmann & Frey, 2016a). From permafrost soils, bacterial candidate phyla OD1, TM7, GN02 and OP11 forming the superphylum Patescibacteria were recovered besides well-established phyla, such as Proteobacteria, Verrucomicrobiota and Acidobacteriota and were found to represent one third of the entire community (Frey *et al*., 2016). At lower elevations, e.g. in alpine grasslands, bacterial communities are primarily dominated by Acidobacteriota (subgroup6, Acidobacteria), Actinobacteriota (Actinobacteria, Thermoleophilia), Proteobacteria (Alpha- and Gammaproteobacteria), Bacteroidota (Bacteroidia), and Verrucomicrobiota (Yuan *et al*., 2014, 2015; Yashiro *et al*., 2016; Chen *et al*., 2020, 2021; Ji *et al*., 2020). Gemmatimonadota and Bacillota (formerly Firmicutes) are further phyla that are commonly present in alpine grasslands, such as in the Tibetan Plateau (Jiang *et al*., 2021). Studies specifically addressing archaeal communities in grasslands are still lacking. However, existing work suggests that Thaumarchaeota (Nitrosphaeria), Nanoarchaeota (Woesearchaeota), Crenaerchaeota (Bathyarchaeota, Thermoprotei), and Euryarchaeota (Thermoplasmata, Methanobacteria) represent the most prevalent archaeal phyla (Malard *et al*., 2022). However, ongoing changes in microbial taxonomy, for archaea (and bacteria), facilitated by the widespread availability of genome sequences that led to the development of comprehensive sequence-based taxonomies like the Genome Taxonomy Database (GTDB) (Parks *et al*., 2018) should be considered. Besides, the coverage of archaea can be less than 25% by using universal primers for 16S metabarcoding (Bahram *et al*., 2019) and selective and specific detection of archaea has been rarely done so far and is thus urgently needed. The dominance of Actinobacteriota and Acidobacteriota and especially of ammonium-oxidising archaea is due to their adaptation to the N- and P-limited conditions typical for alpine grassland soils or higher elevation soils (Liu *et al*., 2017; Ma *et al*., 2019; Praeg *et al*., 2019).

Wang *et al*. (2015) reported no clear trend across a transect spanning 3,106 to 4,479 m on Mount Shegyla (Transhimalaya), Tibetan Plateau in the abundances of bacteria and archaea; however, they found that the ratio of bacterial to archaeal gene copy numbers (as a function of abundances) decreased with increasing elevation, highlighting a switch in favour of archaea. Liang *et al*. (2023) compared variations in taxonomic and functional (N cycle) dis/similarity of bacteria across the Tibetan plateau and found that both were more driven by soil abiotic characteristics than vegetation, but with different environmental drivers prevailing for each. Lazzaro *et al*. (2015) observed the lowest bacterial and fungal abundances at the highest site of an elevational transect in the Swiss Uri Alps (1,930–2,519 m, European Alps). Similar to findings for phylogenetic marker genes, functional gene abundance and diversity were shown to vary with elevation in these studies. Yang *et al*. (2014), in turn, studied the functional diversity at four sites along an elevational gradient in the Qilian Mountains (Tibetan Plateau). Abundance of the Rubisco (ribulose-1,5-bisphosphate carboxylase/oxygenase, involved in CO2-fixation) gene was lower at the lowest site compared to the other sites, which might indicate lower CO2-fixation activities (Yang *et al*., 2014; Guo *et al*., 2015). A succession of the functional genetic potential has also been demonstrated in Swiss glacier forefields (Feng *et al*., 2023). Given that the majority (approximately 99%) is not cultivable, high-throughput sequencing (HTS) has become a powerful tool for assessing and comparing the diversity of prokaryotes, and metagenome assembled genomes increasingly help to describe the uncultivable majority (Hug *et al*., 2016). In alpine systems, the composition, distribution, and structure of microbial communities depend on a number of environmental factors such as temperature, precipitation, and other climatic variables such as moisture and snow cover duration (Malard *et al*., 2022), substrate and nutrient availability, biotic interactions, slope aspect (Adamczyk *et al*., 2019), as well as soil physicochemical and vegetation properties (Shen *et al*., 2015; Donhauser & Frey, 2018; Adamczyk *et al*., 2019; Praeg *et al*., 2019, 2020; Liang *et al*., 2023). While temperature and precipitation typically have a direct effect on microbial communities, the effects of soil and vegetation properties are likely indirect and depend on climatic variables as well as biological and chemical feedback. Yet, the establishment of plants has been identified as an important driver of prokaryotic community structure during early succession (Rime *et al*., 2015; Wojcik *et al*., 2020). Overall, edaphic factors such as soil pH, organic matter content, water, and available P concentrations (Yashiro *et al*., 2016; Bueno de Mesquita *et al*., 2020) remain the main determinants of bacterial and archaeal richness, diversity, and community composition. Soil transplantation experiments to study the changes in the taxonomic and functional gene structures of microbial communities with warming (Zumsteg *et al*., 2013a; Rui *et al*., 2015) confirm field observations. In these studies, changes in the structure of the community were attributed to temperature, moisture, soil properties, and vegetation parameters. By condensing information on community composition to microbial richness and diversity indices, it was shown that bacterial richness decreased with increasing elevation (Shen *et al*., 2015; Adamczyk *et al*., 2019; Praeg *et al*., 2019), whereas for archaea, Singh *et al*.(2012) documented a peak in alpha-diversity at mid-elevations along a 1,000–3,760 m gradient on Mount Fuji (Kantō Mountains), Japan. In glacier forefield soils, microbial community composition was reported to shift in response to increasing C content in soils, decreasing soil pH, and plant establishment (Zumsteg *et al*., 2012). Temperature further affects microbial communities when it reaches extremes (>25 °C) and passes a tipping point where microorganisms react to further temperature increase with pronounced non-linear responses in community-level growth rates, changes in the temperature sensitivity of bacterial growth (Q10), and alterations in community structure (Donhauser *et al*., 2020, 2021). While fungal communities are tightly associated with plants (see Section V.2), bacterial and archaeal communities are also influenced by other prokaryote communities (Malard *et al*., 2022). Soil functions and processes are driven by microbial interactions, and the study of network interactions among bacterial, archaeal, and fungal microbiota is gaining interest. In alpine grasslands, soil pH was found to be a key driver for predicting network-level topological features of soil microbial co-occurrence networks; with increasing soil pH, associations between microorganisms were enhanced and networks became more stable (Chen *et al*., 2021).

Overall, central gaps in knowledge about prokaryotic diversity in alpine soil exist, firstly, in a geographical context. Considering the data from Fig. 4, it is obvious that the global distribution of microbial studies is heterogeneous. Secondly, a deficit we observe is the lack of knowledge about the activity and ecological functions of prokaryotes in situ. While molecular data provide information about phylogeny, conclusions about the function of specific clades are often drawn from few cultured isolates which may not be representative for the entire group (e.g. Verrucomicrobiota). Furthermore, there is a growing need to increasingly take into account the complex interactions among microorganisms themselves, as well as their relationships with plants and soil organisms, when investigating and evaluating microbial diversity.

### (2) Fungi

Fungi comprise a large ecologically heterogenous group of microorganisms (Stajich *et al*., 2009), of which only 2–6% of the estimated 1.5–12 million species have been formally described (Taylor *et al*., 2014; Hawksworth & Lücking, 2017; Bhunjun *et al*., 2022). Functionally, fungi range from ecosystem recyclers, as the main saprotrophic decomposers of (recalcitrant) organic materials (Baldrian & Valásková, 2008; Finlay & Thorn, 2019), to those forming a great diversity of symbiotic associations with plants and animals (Mueller & Gerardo, 2002; Crowther, Boddy & Hefin Jones, 2012; Genre *et al*., 2020). It is this multifunctionality that makes fungi essential components of soil biodiversity in all terrestrial habitats (Wagg *et al*., 2014), including high alpine ecosystems.

To date, much of what is known concerning alpine soil fungal communities comes from European and North American studies of the macroscopic reproductive structures (the sporocarps) produced by fungi. Studies include taxonomic works (Horak, 1993; Cripps, Larsson & Horak, 2010), community and biogeographic studies (Senn-Irlet, 1988, 1993; Ronikier, 2008), and ecological investigations (Graf, 1994). These studies have identified a rich diversity of saprotrophic and plant associated symbionts, many of which appear to be restricted to alpine and arctic environments (Cripps *et al*., 2019).

There are comparably few metabarcoding studies that have included analyses of whole fungal communities in alpine soils, although elevational gradient studies often comprise samples from vegetation neighbouring high-elevation treelines, including alpine heaths (e.g. Tonjer *et al*., 2021). Alpine grassland fungal communities have been investigated using metabarcoding approaches in China (Yang *et al*., 2017; Jiang *et al*., 2018; Zhang *et al*., 2020; Zhang & Fu, 2021) and Central Europe (e.g. Pellissier *et al.,* 2014; Praeg *et al.,* 2019), regions which comprise 79% of the fungal diversity studies found (Fig. 4). These studies demonstrate that different functional groups show a range of responses to changes in elevation, temperature, N addition, and grazing management. In the European Alps, fungi are locally diverse (Brunner *et al*., 2017; Adamczyk *et al*., 2019; Praeg *et al*., 2019; Arraiano-Castilho *et al*., 2021), similar to other alpine regions (Bjorbækmo *et al*., 2010; Perez-Mon, Frey & Frossard, 2020; Rüthi *et al*., 2020). In alpine grassland, soil fungal communities are primarily composed of Ascomycota and Basidiomycota, but also comprise large proportions of unidentified fungi (Pellissier *et al*., 2014; Malard & Pearce, 2018; Praeg *et al*., 2020). Within these phyla, Agaricomycetes (Basidiomycota), Archaeorhizomycetes (Ascomycota), Sordariomycetes (Ascomycota) and Leotiomycetes (Ascomycota) are the most abundant classes of fungi in grasslands (Pellissier *et al*., 2014; Pinto-Figueroa *et al*., 2019). Agaricomycetes are commonly saprotrophic (decomposers) and actively participate in the decomposition of organic matter (Ludley & Robinson, 2008; Edwards & Zak, 2010), especially in cold and dry environments (Ludley & Robinson, 2008). Sordariomycetes and Leotiomycetes are ecologically diverse and include pathogens of either plants and animals, mycorrhiza and plant endophytes, as well as saprotrophs (Maharachchikumbura *et al*., 2016; Johnston *et al*., 2019). Finally, the Archaeorhizomycetes are a widely distributed and abundant class of terrestrial fungi, yet, their role in the ecosystem is still debated (Rosling, Timling & Taylor, 2013; Pinto-Figueroa *et al*., 2019) and their detection is hampered by using the ITS2 region instead of the 18S rRNA gene (Tonjer *et al*., 2021). Archaeorhizomycetes are believed to be associated with plant roots, but experiments suggest they are neither mycorrhizal nor pathogenic (Rosling *et al*., 2013).

Fungal community composition has been intensively studied in glacier forefields where it was shown that the active fungal community composition changes according to soil developmental stages (Zumsteg *et al*., 2012, 2013b; Rime *et al*., 2015; Sannino *et al*., 2020). The diversity of fungi was surprisingly high in barren ground closest to the glacier tongue and was similar to older vegetated soils (Rime *et al*., 2015; Dresch *et al*., 2019). Glacier ice is considered as a fungal inoculum source for the earliest ice-related barren ground and for later plant-covered soil (Rime *et al*., 2016a). Besides the glacier environment, permafrost soils also provide living space for numerous fungi from the prehistoric era (Frey *et al*., 2016; Luláková *et al*., 2019; Pontes *et al*., 2020; Frey, 2021). European permafrost soils are dominated by lichenised fungi and basidiomycetous *Rhodotorula*, including the genera *Naganishia, Mrakia*, or *Leucosporidium* (Frey *et al*., 2016; Adamczyk *et al*., 2021; Sannino *et al*., 2021).

Historically, below-ground studies of alpine fungi have focused on ectomycorrhizal (EM) and arbuscular mycorrhizal fungi as well as root associated symbionts of plants. However, there are very few studies on fungal symbionts associated with ericaceous plants, despite the importance of heath vegetation in alpine systems (Kivlin *et al*., 2017). EM fungi are nevertheless essential for establishment and habitat colonisation by alpine plants such as willows (Nara & Hogetsu, 2004). EM fungi have been examined on a range of hosts, using combinations of linking sporocarps to associated EM tips (*Salix herbacea* L., Graf & Brunner, 1996) or selection of EM tips followed by molecular identification (*Dryas* sp. and *Salix* sp. (Kernaghan & Harper, 2001); *Arctostaphylos uva-ursi* (L.) SPRENG. (Krpata *et al*., 2007); *Bistorta vivipara* (L.) DELARBRE (Thoen *et al*., 2019)). Gao & Yang (2016) used a cloning approach to examine mycorrhizal fungi on herbaceous plant roots in alpine meadows in Southwestern China. However, metabarcoding studies have provided more comprehensive assessments of root associated fungi on particular host species, including *Arctostaphylos* sp. (Hesling & Taylor, 2013), *Dryas* sp. (Bjorbækmo *et al*., 2010), *Carex myosuroides* VILL. (Mühlmann & Peintner, 2008), *Bistorta vivipara* (Mühlmann, Bacher & Peintner, 2008), *Salix* spp. (Ryberg, Andreasen & Björk, 2011). The recent barcoding study by Arraiano-Castilho *et al*. (2021) demonstrated that habitat was a stronger determinant than the host plant for EM fungal distribution in alpine habitats. All of these studies highlight the high diversity of fungal symbionts supporting alpine plants.

Fungi carry out a multitude of functions in ecosystems, and although they interact with many trophic groups, the major focus has so far been on plant associated symbionts. The importance of fungal/plant interactions in the development of plant communities has been particularly well investigated at glacial fronts in alpine zones using metabarcoding studies in Norway (Blaalid *et al*., 2012), the Central European Alps (Brunner *et al*., 2011; Rime *et al*., 2015), and North America (Jumpponen *et al*., 2015).

Fungal communities in alpine grasslands are primarily affected by edaphic, climatic, and biotic parameters. Specifically, soil pH, soil organic C, N, P, soil water content and electrical conductivity are important soil variables, with snow cover duration also exerting an important influence on fungal richness (Pellissier *et al*., 2014; Yang *et al*., 2017; Malard *et al*., 2022). The importance of microtopography in alpine zones, particularly differences in snow lie, is widely recognised in structuring plant communities (e.g. Carlson *et al*., 2015), and a number of studies have also shown topography to be an important driver of soil fungal community composition (Zinger *et al*., 2009, 2011; Frey *et al*., 2016). However, this importance is confounded by the close vegetation/fungal relationships. Further studies on individual plant species (see (Yao *et al*., 2013) over a range of topographies may provide greater insights into the direct role of soil conditions on structuring communities.

The strong connections and dependencies between above-ground plant and below-ground fungal communities (see Yao *et al*., 2013; Tonjer *et al*., 2021) illustrate that climatic and pollutant-induced changes in alpine plant communities (see Steinbauer *et al*., 2018) will have major impacts on the associated soil fungi. The upwards migration of treelines (Harsch *et al*., 2009; Bryn & Potthoff, 2018) and expansion of trees and shrubs into formerly grazed areas (Dibari *et al*., 2020) will, in particular, have significant impacts on both the taxonomic and functional attributes of alpine soil fungal communities. Similarly, the invasion of alien weed species into alpine vegetation, although currently still limited (Alexander *et al*., 2016), could lead to alterations of the indigenous fungal communities (Johnston & Pickering, 2001). Lastly, plant richness and diversity are key to fungal alpha and beta diversity in alpine grasslands (Pellissier *et al*., 2014; Yang *et al*., 2017; Malard *et al*., 2022). Similarly, elevated nitrogen deposition induces major shifts in soil fungal functional groups (van der Linde *et al*., 2018; Zhang, Chen & Ruan, 2018). Coupled with these effects of vegetation change and nutrient availability, there are also direct impacts of changing environmental conditions on fungal communities, with both temperature and moisture being strong drivers of community structure at local (Yao *et al*., 2013), regional (van der Linde *et al*., 2018), and global scales (Tedersoo *et al*., 2014).

### (3) Protists

Protists are defined as all eukaryotes that are not plants, metazoans, or fungi (O’Malley, Simpson & Roger, 2013). They form a vast paraphyletic entity spanning the whole eukaryotic tree of life, comprising large, phylogenetically and functionally diverse groups, and are represented mainly by microbial unicellular organisms (Adl *et al*., 2019; Burki *et al*., 2020).

Due to their phylogenetic, morphological, and functional diversity, it is difficult to generalise findings on the entire protist community. The total diversity of protists in general is unknown, most species are undescribed, and their distribution and functions are poorly understood. Accordingly, knowledge on soil protists is lagging behind that of many other soil organisms (Geisen *et al*., 2018; Bonkowski, Dumack & Fiore-Donno, 2019), and this is reflected in the number of studies focusing on protist diversity in alpine soils (Fig. 4, Table S5). Due to the methodological challenges associated with the study of soil microorganisms, many of which cannot be easily grown in the laboratory, the diversity of protists living in oceans and freshwater ecosystems is better documented than that of soil protists. However, high throughput sequencing studies are revealing that their diversity is highest in soils, partly due to the strong heterogeneity of the soil environment and diversity of soil types (Singer *et al*., 2021). Thus, while numerous studies have explored the diversity of individual protist groups in mountain soils, documentation of this diversity beyond high throughput sequencing approaches still represents a largely open field of research, as is true for minute and microscopic soil organisms in general (Decaëns, 2010). While some protist taxa were described in the European Alps, such as the family Grossglockneriidae (Petz *et al*., 1986; Foissner, 1999), and *Puytoracia jenswendti* SANTIBÁÑEZ ET AL. 2011, a euglyphid testate amoeba discovered on glaciers in the Andes (Santibáñez *et al*., 2011), the true degree of endemicity among alpine protist taxa remains to be determined (Ronikier & Ronikier, 2009).

Diversity patterns of soil protists along elevation gradients have primarily been investigated for specific groups, such as testate amoebae, a group of shelled protists commonly used as models for biogeographic studies. Along such gradients, contrasting patterns of distribution were observed: a hump-shaped pattern along the gradient (e.g. Krashevska *et al*., 2007; Krashevska, Maraun & Scheu, 2010; Lamentowicz *et al*., 2013), the lowest diversity at mid-elevations (Tsyganov *et al*., 2022), decreasing richness, diversity, and equitability with increasing elevation (Heger *et al*., 2016), or no response to elevation (Mitchell, Bragazza & Gerdol, 2004).

With the rise of the molecular era, it has become possible to study the response of the whole community, and Shen *et al*. (2014) showed that an elevational gradient induced little shaping force on protistan communities, which were more strongly influenced by edaphic factors such as soil pH. These contradictory patterns reflect the high diversity of protists, but also likely the fact that some groups are poorly recovered, either due to the fact that primers are not totally universal (e.g. Amoebozoa are typically under-estimated) or that the barcode used (e.g. V4 region of the SSUrRNA gene) contains insertions (e.g. some common soil Rhizaria) that make it impossible to use short reads, as in Illumina sequencing (Pawlowski *et al*., 2012).

Protists are paraphyletic and comprise microeukaryotes that are similar in size and shape to yeasts, but also comprise taxa that are several millimetres in size or even reach several decimetres, such as e.g. slime moulds. It is therefore not surprising that protists do not necessarily respond uniformly to environmental gradients. Nonetheless, as in other habitats, the majority of protistan taxa in alpine soils are believed to be small, motile, and cyst forming bacterivores (Oliverio *et al*., 2020; Kang *et al*., 2022). Accordingly, only a small effect of elevation on alpha and beta diversities of protistan communities can be expected. Stramenopiles, Alveolates, and Rhizaria (SAR), along with Amoebozoa and Archaeplastida dominate the protist diversity in alpine grasslands (Seppey *et al*., 2020). In terms of function, consumers are followed by parasites and phototrophs (Mazel *et al*., 2022). The dominance of consumers in mountain open habitat soils suggests that this functional group could be key in the cycling and turnover of nutrients in this type of ecosystem (Geisen *et al*., 2015; Oliverio *et al*., 2020).

As it is known from low-elevation soils, edaphic factors (e.g. soil moisture, C content, and soil pH) and the local plant community are strongly determining factors of soil protist communities (Oliverio *et al*., 2020; Aslani *et al*., 2022). Besides edaphic factors, temperature and slope of mountain systems also drive protist community assemblages (Seppey *et al*., 2020; Malard *et al*., 2022). Likewise, Hu *et al*. (2022) showed a strong influence of soil moisture and N content as shaping factors of soil protistan communities at high elevations, while Shen *et al*. (2014) found protist communities to be primarily correlated with soil pH. Only a few studies have aimed at inventorying protist communities in alpine ecosystems (Hu *et al*., 2022; Kang *et al*., 2022). These studies claim that body size determines the assembly of protist communities, with deterministic factors (e.g. soil acidity, temperature) being more important in protists than in other microbes. Furthermore, Kang *et al*. (2022) showed that the turnover rates among alpine environments were lower in protists than in other microorganisms (bacteria and fungi), which they explain with a higher dispersal rate of motile protists. However, this finding contrasts with floodplains where protists showed higher spatial patterns while bacteria communities changed primarily seasonally (Fournier *et al*., 2020). Borg Dahl *et al*. (2019) highlighted the importance of plant community as a major determinant of myxomycetes in the European Alps. As such, physicochemical properties and vegetation patterns will differentially shape protists in alpine forests, shrublands, grasslands, pastures, and high alpine zones. To test these suggested trends and hypotheses on protist communities, more targeted inventories are needed across alpine systems. A recent study manipulated precipitation, warming, and nitrogen addition in alpine habitats, revealing that these global change factors fundamentally alter soil protist communities and their abundances. In this study, precipitation and nitrogen input caused an increase in protist diversity and abundance, respectively, while decreased precipitation and warming reduced them (Hu *et al*., 2022). Further, it can be expected that changes in bacterial, fungal, but also plant and animal communities will cascade to protists (Valencia *et al*., 2018). Therefore, climate change is expected to alter protist communities in alpine habitats with potential impacts on other components of the soil microbiome and on soil functions (Mazel *et al*., 2022).

The last decades have revolutionised the perspective on soil protist functional roles, which span the whole spectrum from predators, primary producers, parasites, decomposers, phototrophs and saprotrophs (Geisen *et al*., 2018, 2020). In soils, protists feed on a wide variety of substrates, with heterotrophs representing the most abundant and diverse functional group (Bonkowski *et al*., 2019). Soil protists were first shown to be key bacterial predators that control bacterial abundances and, via the microbial loop, make nutrients available for plant growth (Clarholm, 1985). However, protist predators occupy different trophic niches by feeding on other microorganisms like bacteria, fungi, algae, micro-metazoa such as nematodes and rotifers (Yeates & Foissner, 1995; Gilbert *et al*., 2000; Jassey *et al*., 2013; Geisen *et al*., 2015; Estermann *et al*., 2023), and other protists (Seppey *et al*., 2017; Geisen *et al*., 2018; Bonkowski *et al*., 2019). Hereby, phagocytosis appears to be the main mechanism for nutrient acquisition (Singer *et al*., 2021).

Bacterivorous taxa may dominate the protist community in many cases (Oliverio *et al*., 2020; Aslani *et al*., 2022), but the dominant feeding habits can be expected to match the available resources and especially the bacteria to fungi ratio, which responds to soil pH (Rousk, Brookes & Bååth, 2009). Hence, fungivores are likely more common in subalpine (e.g. conifer-dominated forests) and lower alpine (e.g. ericoid heath) habitats, as perfectly illustrated by the obligate fungivorous grossglocknerid ciliates that were discovered in the European Alps (Petz *et al*., 1986; Foissner, 1999).

Soil protists, including crop pathogens like *Phytophthora infestans* (MONT.) DE BARY, have a broader role as parasites, potentially affecting plant or soil animals. However, the diversity and interactions of these protist parasites remain understudied. Apicomplexa parasites of invertebrates and vertebrates, strongly dominate soil protist diversity in tropical forests and this dominance is thought to reflect the overall invertebrate diversity throughout the ecosystem, from soil to canopy (Mahé *et al*., 2017). In line with this, a study in the Swiss Alps showed that the diversity of Apicomplexa in various alpine habitats correlated positively with the diversity of their putative metazoan hosts (Singer *et al*., 2020). The relative contribution of parasites to the total protist community compared to other functional groups was, however, shown to decrease with increasing elevation, likely due to the reduction in host density with elevation (Mazel *et al*., 2022).

Phototrophic protists, like *Chlorella* and *Trebouxia*, are common as symbionts in lichens, but also as free-living forms at the soil surface (Jassey *et al*., 2022). However, the abundance of free-living phototrophic protists (and their predators) is highest in moist (e.g. peatlands) and open (e.g. arid or alpine) vegetation (Gilbert *et al*., 1998; Seppey *et al*., 2017). In arid habitats, including patchy alpine vegetation, phototrophic protists contribute to the formation of biocrusts, which are major contributors to organic C and N fixation (Dickson, 2000) and reduce soil erosion (Evans & Johansen, 1999). While soil protists have long been neglected in soil microbiological studies (Geisen *et al*., 2020), they now are the focus of an increasing number of studies as their importance as determinants of plant performance is established (Bonkowski, 2004; Gao *et al*., 2019). Thus, protists are now recognised as important elements in soils ecosystems due to their role in the microbial food web and nutrient cycling (Adl & Gupta, 2006; Geisen *et al*., 2016) and their contribution to biogeochemical cycles, especially C (Geisen *et al*., 2020) and silica (Aoki, Hoshino & Matsubara, 2007).

## V. SOIL INVERTEBRATES

#### KEY ASPECTS

- The treeline ecotone harbours a high diversity of soil macro-, meso-, and micro-invertebrates, as species from both forest and grassland ecosystems coexist. At higher elevation, the shallow soils are mainly inhabited by soil meso- and micro-invertebrates.
- Faunal diversity generally decreases with increasing elevation, as climatic and energetic conditions become more challenging. Some taxa reach their upper distribution limits (e.g. earthworms and millipedes in the high alpine zone).
- Essential ecosystem functions are carried out by only a few key taxa (e.g. litter decomposition in the high alpine is mainly carried out by Nematocera larvae and soil meso-invertebrates).
- Soil food webs in high alpine soils are simple, with fewer interactions compared to lowland soils. Omnivorous and opportunistic feeding habits have increased to ensure energy intake.
- Extensive grazing by livestock and wild ungulates can improve conditions for soil fauna by providing nutritious manure and reducing cover of recalcitrant dwarf shrubs.
- We found 205 publications dealing primarily with alpine soil fauna (i.e. 118 for macro-, 52 for meso-, and 35 for micro-invertebrates), mainly from the mountain regions of Central & Southern Europe (42.9%), Central Asia (15.6%), and Australia & New Zealand (13.2%); see Fig. 5 and Table S6.

In general, soil invertebrates belong to a wide range of taxa. Their diversity is particularly high close to the treeline, as representatives of all size classes (micro-, meso-, and macro-invertebrates, (Orgiazzi *et al*., 2016) are found and grassland species co-occur with forest species, especially in intertwined dwarf shrub habitats. Macro-invertebrates (taxa with a mean body size > 2 mm, mainly earthworms, spiders, myriapods, isopods, ants, and insect larvae) decrease in numbers with increasing elevation due to climatic and topographic factors. Vegetation cover and the amount of soil decrease, particularly limiting soil macro-invertebrates that require a vegetation cover that produces litter material and/or the physical habitat space that is provided by mature soils. Meso-(mainly collembolans, mites, and enchytraeids) and micro-invertebrates (mainly nematodes, rotifers, and tardigrades) can still be abundant in shallow high-elevation soils even though habitat space and litter inputs from vegetation are reduced.

Compared to soil microbiota, data on soil fauna are scarce and often limited to ground-dwelling taxa (Burton *et al*., 2022). Soil fauna studies focusing on alpine and high-elevation habitats are especially rare. Available data pertain primarily to alpine regions in Central & Southern Europe (i.e. 42.9%, Fig. 4 and Table S6) such as the Central European Alps (e.g. Puntscher, 1980; Meyer & Thaler, 1995; Koch & Erschbamer, 2010; Gobbi *et al*., 2020; Seeber *et al*., 2021), Central Asia (15.6%) such as the Tibetan Plateau (e.g. Wu, Zhang & Wang, 2015; Devetter *et al*., 2017), and Australia and New Zealand (13.2%, e.g. Salmon, 1940; Hammer, Foged & Nørvang, 1966; Houston & Greenslade, 1994; Minor *et al*., 2016; Mesibov, 2018; Green & Slatyer, 2020), while alpine regions in the Americas (i.e. Rocky Mountains, Appalachians, and Andes), Africa (i.e. Drakensberg), and the Caucasus remain understudied (but see Armstrong & Brand, 2012; Kokhia & Golovatch, 2020). We were able to find only five soil fauna publications for each of these alpine regions (Table S6, Appendix S1). A similar outcome was found for the alpine region of Siberia (i.e. North Asia), where information is locally available but mainly published in Russian and therefore not indexed in the ‘Web of Science’ portal.

In alpine environments, large soil fauna is generally sampled by installing pitfall traps (macro-, partially also meso-invertebrates), by taking soil core samples (all groups), by suction sampling (ground-dwelling meso-invertebrates), as well as via hand sorting and hand sampling (macro-invertebrates); pitfall traps are preferably used in higher elevations as soil is getting scarce and shallow. To cope with methodological and logistical limitations, additional approaches such as soil biodiversity indices (e.g. QBS-ar, Maienza *et al*., 2022) and DNA metabarcoding (e.g. via environmental DNA (eDNA), Rota *et al*., 2020) are increasingly applied also in alpine habitats. Amongst the alpine soil fauna species described to date, some are rarely found and observations are often based on occasional records. Such observations can even lead to new discoveries for alpine regions due to the scarcity of research such as the carabid beetle *Orthoglymma wangapeka* LIEBHERR, MARRIS, EMBERSON & SYRETT & ROIG-JUÑENT, 2011 (Liebherr *et al*., 2011) and the oribatid mite *Crotonia ramsayi* COLLOFF, 2015 (Colloff, 2015) for New Zealand, the isotomid springtail *Skadisotoma inpericulosa* GREENSLADE & FJELLBERG, 2015 (Greenslade & Fjellberg, 2015) for Australia, and *Opetiopalpus sabulosus* MOTSCHULSKY, 1840 (Steinwandter *et al*., 2019). A high percentage are regionally endemic or found in restricted geographical areas, as observed in the European Alps (e.g. Komposch, 2011), in Australasia (Boyer & Giribet, 2009), and in the Drakensberg of Southern Africa (Armstrong & Brand, 2012). These species are mainly relicts of the last glaciations that survived in nunataks and other refugia offered by highly heterogeneous mountain topography (Brighenti *et al*., 2021); in subsequent interglacial periods, these alpine invertebrates have expanded extensively (Hill *et al*., 2009). For Australasian alpine taxa, a deeper phylogeographic structuring was shown compared to European and North American ones, possibly reflecting less intense glaciation and a higher availability of refuges during glaciation events (King *et al*., 2020). Colonisation processes in high alpine areas can be surmised by observing the colonisation of alpine land when glaciers retract (Koch & Kaufmann, 2010; Hågvar *et al*., 2020). The first to re-colonise the bare land, which is comparable to quarries, are mostly agile ground-dwelling predators (e.g. carabid beetles, harvestmen, lycosid spiders) depending presumably on windblown animals as food sources, followed by the meso-invertebrates (springtails and oribatid mites) after 30 years. Finally, larger detritivores (millipedes and Nematocera larvae) appear when the soil and the vegetation are more developed (Kaufmann, Fuchs & Gosterxeier, 2002).

### (1) Macro-invertebrates

The diversity of soil macro-invertebrates – often referred to simply as soil macrofauna – is generally lower in high alpine grasslands than in the alpine and subalpine zone and often peaks in ecotone areas (i.e. the transition zones at the treelines and open alpine grasslands; Fontana *et al*., 2020; Steinwandter & Seeber, 2023). Elevation and vegetation are the primary determinants of alpine soil macro-invertebrate communities (Kooch & Noghre, 2020; Steinwandter *et al.,* 2022; Xie *et al.,* 2022; Lavelle *et al.,* 2022). Earthworms (Lumbricidae), for instance, show a hump-shaped distribution peaking at the treeline ecotone area (Fontana *et al*., 2020; Gabriac *et al*., 2023). Earthworm abundance decreases in alpine grasslands (Seeber *et al*., 2005; Steinwandter *et al*., 2018), likely because of their limited tolerance to the cold temperatures encountered at higher elevations (Meshcheryakova & Berman, 2014). Additional influencing factors include vegetation attributes such as plant life-forms and host-plant distributions (e.g. Edwards & Arancon, 2022), as well as soil attributes such as pH, clay, and water content. Also, poorly developed soils provide limited habitat space for burrowing species. Yet, abundances and species diversity may increase with the presence of grazing livestock and wild mammals, whose dung represents a readily available food source for all decomposer taxa (Bueno & Jiménez, 2014; Steinwandter *et al*., 2018; Jászayová *et al*., 2023).

Millipedes (Diplopoda) are litter-dwellers and therefore mostly found in dwarf shrub-rich grasslands and ecotones above the treeline, where they find the mature and stable soils they prefer as well as more abundant food resources such as litter and organic debris (Onipchenko & Zhakova, 1997; Steinwandter *et al*., 2018; Gobbi *et al*., 2020; Kokhia & Golovatch, 2020). Most millipede species reach their upper limit of distribution at the transition between the subalpine and alpine zones and are rare or even completely absent in high alpine habitats. Soil core samples from high elevations generally contain few to no millipede specimens, making estimates of their densities difficult. However, millipedes (and soil invertebrates in general) are easier and more efficiently detected in high alpine environments by using pitfall traps or hand sampling and are therefore occasionally found at higher elevations (e.g. *Beronodesmoides* spp. (Polydesmida: Paradoxosomatidae) up to 4500 m in Nepal (Golovatch, 2015). Millipede species inhabiting high elevations mainly belong to the orders Polydesmida, Chordameutida, and Julidae (Beron, 2008, 2016). In the Central European Alps, species found in higher elevations belong mainly to Chordeumatida, which are described to be petrophilic with a preference for cold mountain areas and which are active beneath the snow (Meyer, 1980). These millipedes were found in high numbers at sites up to 3000 m (Steinwandter & Seeber, 2023), while other myriapods such as centipedes (Chilopoda) were almost absent. Other millipede species that frequently inhabit European mountain soils and can be found at high elevation include eurytope millipede species such as *Ommatoiulus sabulosus* (LINNAEUS, 1758) (Julida: Julidae) as well as specialists such as the endemic *Glomeris transalpina* KOCH C. L., 1836 (Glomerida: Glomeridae); both are known to inhabit alpine rocky sites and soils even up to 3000 m. However, elevational limits may now change with ongoing global warming: Gilgado *et al*. (2021) recently described ten millipede species whose elevational limits in the Swiss Alps have expanded upwards by several 100 metres over the last century.

Surface-active and highly mobile and agile predators, such as spiders, harvestmen, and some beetle families are abundant representatives of the high alpine soil fauna (Kaufmann *et al*., 2002; Hågvar *et al*., 2020; Gilgado *et al*., 2022) and seem not to depend on mature soils but rather on available prey. Numerous studies have investigated the diversity of beetles in mountain soils, but the majority focus on a few widely distributed and well-known families (e.g. Carabidae, Staphylinidae, and Scarabaeidae). The density, diversity, and distribution of predatory beetles are affected by a wide range of factors such as biotic interactions, vegetation (Negro *et al*., 2010; Yu *et al*., 2013), abiotic factors such as temperature and moisture (Yu *et al*., 2013), historical factors such as climatic variability and topographical changes, and human activities (Larsen, 2012; Brandmayr & Pizzolotto, 2016). Further, topographic isolation may boost beetle diversity as it was found by Armstrong & Brand (2012) on isolated peaks (i.e. > 3000 m) of the Drakensberg (Afro-alpine Region), where leaf-(Chrysomelidae), ground-(Carabidae), and sap beetles (Nitidulidae) dominated.

In the case of ants (Hymenoptera: Formicidae), as with other soil fauna, their presence and abundance decrease with increasing elevation in alpine settings, where severe filtering can be detected both on taxonomic as well as functional and phylogenetic diversity (Glaser, 2006; Machac *et al*., 2011; Chaladze, 2012; Reymond *et al*., 2013; Bishop *et al*., 2014). Elevational limits to occurrence are related to the ability of ants to cope with low temperatures (Bishop *et al*., 2017). However, while ant colonies tend to occur in the lower alpine area and become increasingly absent in high alpine areas, some individuals (mainly specimens from the winged ‘alates’ caste) may be transported upwards by wind. Overall, ant diversity peaks at mid-elevation and decreases constantly – and often linearly – with increasing elevation (Subedi & Budha, 2020). Similar results were found for the Maloti-Drakensberg in Southern Africa by Bishop *et al.,* (2014) who attributed the spatial and temporal differences primarily to temperature. In the European Alps, most ant species occurring in the alpine habitat are also present in the higher montane forest belt (Glaser, 2006), and a higher species diversity was recorded in the treeline ecotone (Guariento & Fiedler, 2021). Interestingly, a high number of social parasitic ant species are reported from alpine habitats without a clear explanation so far, except that the harsher environment might positively select for such life history traits (Dunn *et al*., 2009; Schifani *et al*., 2021). In alpine grasslands, the effect of ants on soils is mostly related to nest construction, since most taxa build their nests in the soil, causing soil turnover as well as nutrient accumulation and influencing the vegetation (Wang *et al*., 2017; Zhao *et al*., 2020). Ants strategically establish their nests beneath rocks, leveraging the warmth absorption and insulation (McCaffrey & Galen, 2011). Therefore, the rock features (e.g. distribution, shape) represent an important factor for the establishment of ant nests. Studies investigating the functional role of ants indicate higher trophic levels in the alpine environment (Spotti *et al*., 2015; Guariento, Martini & Fiedler, 2018), even suggesting intraspecific dietary shifts (Guariento, Wanek & Fiedler, 2021).

Other insect taxa such as larvae of lower flies (Nematocera) increase in numbers at higher elevations. They can – at least in parts – carry out crucial ecological functions such as litter decomposition and bioturbation which are usually provided by detritivores such as millipedes and earthworms (Meyer & Thaler, 1995; Kitz *et al*., 2015).

### (2) Meso-invertebrates

Soil meso-invertebrate communities – often referred to simply as soil mesofauna – are mainly composed of springtails (Collembola), mites (Acari), other small arthropods, and potworms (Enchytraeidae) (Potapov *et al*., 2022). Of these, springtails are the most widespread and abundant invertebrates, occurring in almost all terrestrial ecosystems (Hopkin, 1997; Deharveng, 2004). They play essential roles in many soil ecosystem processes, such as C and N cycling, soil microstructure formation, and plant litter decomposition. Collembola density and diversity vary significantly with environmental factors and plant community composition, and in the shallow alpine soils they inhabit mostly the litter and upper soil layers (Seeber *et al*., 2021; Xie *et al*., 2022).

In general, the composition and abundance of meso-invertebrate communities in soil are dependent on elevation (Striganova & Rybalov, 2008; Jiang, Yin & Wang, 2015; Khabir *et al*., 2015; Schatz, 2017; Winkler *et al*., 2018), soil properties (van der Merwe *et al*., 2020), the identity of plant species and the variability of vegetation communities (Eo *et al*., 2016; Xie *et al*., 2022), all of which can lead to a high spatial heterogeneity with many local microhabitats. Furthermore, factors related to climate change, such as temperature (Harte, Rawa & Price, 1996; Alatalo *et al*., 2017) and reduced soil water availability (Sylvain *et al*., 2014), as well as (anthropogenic) disturbances may also affect soil mesofauna diversity and communities. Habitat management (Kooch, Shah Piri & Dianati Tilaki, 2021), tourist activity (Meyer, 1993; Kopeszki & Trockner, 1994), cattle trampling and grazing (Hauck *et al*., 2014; Risch *et al*., 2015), fire (Driessen & Kirkpatrick, 2017), soil erosion (Meyer, 1993; van der Merwe *et al*., 2020), and pollution (Rusek, 1993; Visioli *et al*., 2019) all affect the meso-invertebrates living in alpine grasslands, suggesting that environmental filtering is the predominant process shaping soil meso-invertebrate communities (Visioli *et al*., 2019).

In the nival zone, the soil fauna community is composed almost exclusively of meso-invertebrates – springtails, mites, and by region additionally by predatory false scorpions (Pseudoscorpiones), e.g. in European Alps, Carpathian Mountains. Mesofauna distribution is scattered and limited to favourable refugia such as congregations of detritus or cushion plants (Meyer & Thaler, 1995). Some specialist taxa are adapted to snowbeds, which can persist for most of the year (Seeber *et al*., 2021). These species rely on aeolian food sources (i.e. wind-blown debris) or prey on small animals searching for this food; the different taxa are active at different times of the day as extreme environmental conditions restrict their activity (Mann, Edwards & Gara, 1980).

### (3) Micro-invertebrates

Soil micro-invertebrates (taxa < 0.1 mm in size) – simply referred to as soil microfauna – mainly comprise roundworms (Nematoda), rotifers (Rotifera), and water bears (Tardigrada). Little research on these tiny soil invertebrates has been conducted in alpine regions (Devetter *et al*., 2017), however, a global distribution map of nematodes revealed a positive relationship between organic C content in mountain soils and abundance of nematodes (van den Hoogen *et al*., 2019). Micro-invertebrates are generally favoured in fertile soils with high contents of N, P, and organic matter (Devetter *et al*., 2017), and are easily affected by disturbances such as soil degradation and shrub encroachment after abandonment (Hu *et al*., 2017; Wu *et al*., 2017; Wang *et al*., 2018)(Hu *et al*., 2017; Wu *et al*., 2017; Wang *et al*., 2018). Recently, Porazisnka *et al*. (2021) showed that soil nematodes expand their distribution ranges with elevation by following expanding plant species, and as plant communities become more complex and diverse even at higher elevations, a more diverse nematode community may increasingly contribute to C and N sequestration. Further, Li *et al*. (2023) found that soil nematodes – which are generally water-bound – respond positively to higher precipitation and soil water content in alpine grasslands of the Tibetan Plateau, with higher trophic-level nematodes (i.e. omnivores, carnivores) showing stronger effects than lower trophic-level nematodes (i.e. bacterivores, fungivores).

### (4) Adaptation strategies of fauna to mountain soils

The conditions in high alpine areas can be hostile to animal life, but soil taxa have adapted over a long time and have developed strategies to cope with the harsh and varying climate. Many taxa have black, dark brown, or dark grey body colour (e.g. beetles, spiders) (Armstrong & Brand, 2012), an adaptation that allows larger cold-blooded mountain animals to better absorb sunlight and therefore energy. Appendages and legs are often shorter than those of the same species and groups living in lower habitats. Wing size is also often reduced (i.e. a higher degree of brachyptery or winglessness), for example in carabid communities of high alpine habitats (Pizzolotto, 2016). Due to the long duration of snow cover, alpine soil invertebrates show increased cold resistance, behavioural thermoregulation, and actively seek thermally-buffered microhabitats (Dillon *et al.,* 2006; Schoville *et al.,* 2015; Buckley *et al.,* 2015). Yet, individuals often search for food on the snow surface or in more favourable locations that can only be reached by crossing snowfields, thus, the risk of hypothermia is often high.

Beside physiological adaptations, high alpine soil fauna typically show more generalist diets (i.e. omnivory) due to the low availability of food resources. Predation seems to be driven by the presence and abundance of a given prey. For example, in extremely high-elevation environments such as glacier forefields, carabid beetles of *Nebria* spp. and lycosid spiders of *Pardosa* spp. prey on springtails (König, Kaufmann & Scheu, 2011; Sint *et al*., 2019; Hågvar *et al*., 2020), which are specifically tied to the geomorphology of these habitats (i.e. rough stones that can trap food and prevent flushing, Buda *et al*., 2020). Food limitation in such environments results in simpler and more reduced food webs compared to lowland habitats (König *et al*., 2011; Raso *et al*., 2014; Steinwandter *et al*., 2018). Species which are usually saprotrophic also include animal food sources (e.g. exuvia, carcasses, tissue parts) in their diet and may feed on living animal tissue as plant-based litter is rare or even absent. For predators such as carabid beetles living in barren high alpine soils with limited number of prey (i.e. mainly springtails), increased intraguild and intraspecific predation has been observed to sustain nutritional needs (Raso *et al*., 2014). Additional food can also come from airborne sources, including flying and wind-carried insects, as well as detritus (Růžička & Zacharda, 1994; Hågvar *et al*., 2020). However, specialised predation to efficiently intercept the most abundant prey has also evolved, as in the case of the carabid genera *Leistus* and *Notiophilus*, which trap springtails with their antennae setae (i.e. setal traps, Bauer, 1985). Additional adaptations pertain to the invertebrates’ life history. For example, the life cycle of millipedes and other soil invertebrates can be interrupted and postponed to spring of the following year if it cannot be completed within a single season (Meyer, 1985; Sømme & Block, 1991; Valle *et al*., 2020). Also, parthenogenesis is widespread, especially among the soil meso-invertebrates (e.g. springtails and mites), allowing them to thrive when the conditions are more variable (Pan *et al*., 2023).

## VI. KNOWLEDGE GAPS AND RESEARCH OPPORTUNITIES

A number of recent papers discuss data, knowledge, and policy gaps in soil biodiversity science at a global scale (Guerra *et al*., 2020, 2021, 2022), but there has been no such contribution for alpine soils so far. This is the case despite the critical importance of healthy mountain soils for human safety and wellbeing and for the global provisioning of essential goods and services such as clean water. In this section, we address this gap and identify three main fields/topics for future soil biodiversity research and policymaking in mountains. We put a special emphasis on mountain characteristics that are key but also challenging for soil biodiversity research in mountains (see Klein *et al*., 2019). These include the typical elevational gradients encountered in mountains, their remoteness and simultaneous exposure to global change, and their global distribution.

### (1) Opportunity 1: Increase and improve mountain soil biodiversity data

#### Premise

In line with recent global analyses, our synthesis indicates that data on mountain soil biodiversity is generally sparse and biased (Figs. 1 and 3–5) towards specific geographic regions (Central Asia and Central Europe) and taxonomic groups (mostly soil microbiota). In particular, it points to limited data for groups such as soil invertebrates that are more exhaustively described in other biomes (Geisen *et al*., 2017, 2018; Eisenhauer *et al*., 2022). This lack of data on species diversity and occurrence in mountain soils constitutes an important gap in our knowledge of biodiversity on Earth. It causes species to be overlooked by science, conservation, policy, and advocacy, even if they are possibly on the verge of extinction and/or play critical roles in supporting ecosystem functions. This is particularly important in soils, where interactions among taxa are ubiquitous and essential for species persistence and ecological functions (e.g. Bardgett & van der Putten, 2014). As soil organisms and keystone species disappear, the functioning of entire ecosystems could be disrupted, threatening humanity at large (Jousset *et al*., 2017; Banerjee, Schlaeppi & van der Heijden, 2018; Chen *et al*., 2020; Guerra *et al*., 2021).

The observed lack of species data also comes with the risk of overlooking invasive species that could alter soil properties and represent a threat to native species (e.g. the earthworm *Amynthas agrestis* (GOTO & HATAI, 1899) in the Great Smoky Mountains (Appalachian Mountains, Snyder, Callaham & Hendrix, 2011)). In addition to the absence of information on the mere existence of many species, limited (long-term) data on trends in populations, species distributions, and community composition further jeopardise the ability of science and policy to detect, interpret, and ultimately address or prevent effects of global change on mountain soils and ecosystem functions. It equally hinders the detection of potentially unexpected effects of nature conservation. Whereas species distribution and range expansions are increasingly better documented in particular for plants (e.g. Rumpf *et al*., 2018; Staude *et al*., 2022), such information hardly exists for soil biota, and temporal variation in diversity along elevational, topographical or other ecological gradients is largely unknown (e.g. see Seppey *et al*., 2020 for soil protists). Furthermore, limitations in spatial representativeness and coverage of taxonomic groups in soil biodiversity data constrain our capacity to understand mountain soil systems and their response to change based on comparative analyses at multiple biogeographic scales. Given the worldwide occurrence of mountains, the differences between mountain ranges, and the existence of differences even between the south- and north-facing slopes of individual mountains, such comparative approaches are both important and particularly interesting. Importantly, these gaps in knowledge also hamper the establishment, design, and prioritisation of monitoring efforts as well as the integration of soil biodiversity in “Red Lists”.

#### Directions

We support previous calls for a better geographic and taxonomic coverage in soil biodiversity research and for prioritising long-term monitoring of life in mountain soils at national (Guerra *et al*., 2020, 2021; Eisenhauer *et al*., 2022) and global (Maestre & Eisenhauer, 2019) scales. Multiple options exist for increasing and improving mountain soil biodiversity data (see also Hochkirch *et al*., 2021). One resides in molecular approaches such as DNA or RNA barcoding and the use of metagenomics and metatranscriptomics. These methods offer great opportunities, in particular for the detection and identification of microbiota as well as for attributing them to threat categories (e.g. Guerra *et al*., 2021). While in use for bacteria and fungi already (see Section V), they could also deliver much needed information for other soil organisms in mountains. Another option to increase species discovery rates resides in the identification of mountain locations where unknown taxa are most likely to be encountered (e.g. Delgado-Baquerizo, 2019; Verdon *et al*., 2023). Such an effort is particularly interesting in remote locations, where *in situ* sampling is particularly challenging. Similar approaches for the identification of sampling locations based on the intersection of spatial datasets of mountain extents (Snethlage *et al*., 2022), key environmental variables, and abiotic factors (e.g. soil temperature and type) might also serve the prioritisation of appropriate sites for long-term monitoring of mountain soil species and communities along elevational gradients. Furthermore, as suggested by van der Putten *et al*. (2023), an alternative to identifying species is to qualify soil biota based on traits and thereby more readily understand what ecosystem functions are likely to be lost as species go extinct. A trait-based approach could be particularly interesting in mountains where harsh environmental conditions, including extreme temperature (gradients) and biophysical stressors such as recurrent avalanches, are likely determining unique sets of traits.

### (2) Opportunity 2: Increase and improve information on the environmental determinants of biodiversity in mountain soils, the drivers of change in mountain soil biodiversity, and the consequences of these changes

#### Premise

The occurrence and diversity of life forms in mountain soils are strongly determined by environmental factors, including temperature, snow cover, precipitation, humidity and wind, as well as factors such as soil properties (e.g. pH, organic matter quantity and quality, parental material composition). Accordingly, as these factors are changing in response to changes in climate, land-use, and other drivers such as pollution, soil communities are expected to experience novel life conditions influencing their distribution, dynamics, survival potential, and functions (e.g. Feng *et al*., 2023). In that context, glacial forefields represent newly forming ecosystems of particular interest and need of protection (Bosson *et al*., 2023; Tollefson, 2023). Additionally, (expected) change in the elevation range limits, distribution, and community composition of vascular plants in response to global change are likely to have additional consequences on soil organisms, whose ecology and life histories are tightly associated with plants. Furthermore, as environmental factors such as temperature and soil moisture determine not only the occurrence and diversity of organisms but also specific biochemical cycles such as the production of methane (e.g. Hofmann, Reitschuler & Illmer, 2013), feedback loops are likely to magnify effects of climate and land-use change and thereby exacerbate the exposure of soil biota to unprecedented and extreme environments. Such feedback loops or cascading effects are further exacerbated by the reciprocal effects of organisms on their environment (e.g. pedogenic effects of cryptogams, Musielok *et al*., 2021).

#### Directions

Identifying the impacts of global change and anthropogenic activities on mountain soil biodiversity is essential to safeguard soil functions, services, and health (Arora, 2023). In that context, considerable improvements are needed in the spatial resolution, temporal coverage, and accuracy of information on fundamental variables such as soil type, temperature, moisture, pH, and precipitation in mountains (e.g. Randin *et al*., 2020). We also join others (e.g. Bouaicha *et al*., 2022; Eisenhauer *et al*., 2022) in calling for improved data and remote-sensing products on less common drivers of soil biodiversity, such as pollution by microplastics, chemicals, and heavy metals. Such data are particularly important in mountain regions, where global atmospheric transport of micropollutants as well as human activities, such as mining, pastoralism, and tourism are major sources of pollution (Schmeller *et al*., 2022), impacting soils and their biodiversity. Improved data are further important to identify and better understand the interactions of global change drivers, both in space but also in time, as exposure to anthropogenic factors typically varies over the seasons (e.g. pastoralism in the summer and ski runs in the winter). Moreover, given the high level of interactions between soil organisms, which causes conditional dependencies between groups of soil biota and the environment, a holistic approach is needed when making inferences about possible drivers of change or responses to given environmental variables. Accordingly, understanding the response of soil biota to global change calls for the joint monitoring and analysis of multiple groups and species in their interaction with each other and their environment (Eisenhauer *et al*., 2022). Given the worldwide distribution of mountains (Körner *et al*., 2017; Snethlage *et al*., 2022), we further recommend a comparative approach to global change research in mountain soils and take advantage of the fact that mountains across the world differ in their environmental conditions, their history of exposure to human pressure, as well as in their environmental gradients. For example, whereas extreme temperatures are not yet recurrent in most mountain regions worldwide, exposure to high temperatures and extreme dryness is typical in certain ranges such as in the Mediterranean or inner European Alps, where soils and their biota show specific community composition and species adaptations in response to these conditions (Praeg *et al*., 2020). Accordingly, comparative analyses of soil biodiversity, as well as of genetic and trait diversity across mountain ranges are likely to yield interesting understanding with respect to evolutionary potential of terrestrial ecosystem (Bardgett & van der Putten, 2014) in the face of global change. Similarly, beyond the assessment of niche variation along environmental gradients within mountains (e.g. Malard *et al*., 2022), the quantification and comparison of niche properties across environmental gradients in different mountain ranges is expected to help evaluate the potential influence of global change on taxa and communities as well as improving our capacity to predict the fate of ecosystems and thereby inform conservation (e.g. Mod *et al*., 2021). Biogeographic studies and palaeoecological analyses might further provide useful information on the distribution of species over evolutionary times and on the resilience of mountain soils to changes in climate.

### (3) Opportunity 3: Increase policy-relevant mountain soil biodiversity science and improve mountain soil conservation and policies

#### Premise

Belowground biodiversity is essential to healthy soils, which in turn are crucial for food production, aboveground biodiversity, climate control, and human health and security (Banerjee & van der Heijden, 2023). Due to the intrinsic connection between terrestrial and aquatic environments, soil biodiversity and healthy soils are particularly important in mountains in their role as water towers. However, despite their importance and the growing interacting impacts of climate and land-use change, pollution, and overexploitation (e.g. mining) in mountain regions, mountain soils and their biodiversity – even more so than lowland soils – remain only poorly addressed in laws, restoration, and conservation policies (but see Stanchi *et al*., 2023). One of the numerous challenges associated with soil conservation and protection and with formulating laws and guidelines for sustainable use of mountain soils is that soils are connected across national borders and continents by human activity (van der Putten *et al*., 2023), calling for international agreements. An additional difficulty specific to mountains is their transboundary nature, with many mountain ranges crossing national borders, which further requires reinforced international collaboration in the establishment of meaningful policies. In that context, the Soil Conservation Protocol and the Soil Working Group of the Alpine Convention represent valuable efforts. The difficulty of collecting data in mountains further contributes to making their soils and the diversity of species they host a blind spot in science, conservation, and policymaking. The observation that most parties to the Convention for Biological Diversity (CBD) have no national target explicit to soil conservation and biodiversity (Guerra *et al*., 2021) and that the protection and conservation of soil biodiversity and soil ecosystem functioning have been insufficient to date (Zeiss *et al*., 2022) also applies to mountain soils.

#### Direction

We support ongoing efforts (e.g. Guerra *et al*., 2021; Arora, 2023; van der Putten *et al*., 2023) to raise the importance of soil biodiversity in environmental policies and to formulate frameworks for the protection and restoration of soils (e.g. ‘EU Soil Strategy for 2030’ and the associated ‘Soil Monitoring Law’). However, given the critical importance of healthy and biodiverse soils in mountains (e.g. for natural risk regulation), we herewith call for dedicated efforts and explicit political commitments towards their targeted protection. The ongoing development of National Biodiversity Strategies and Action Plans in response to the adoption of the Kunming-Montreal Global Biodiversity Framework represents a unique opportunity to collaborate on the formulation of soil biodiversity conservation targets and policy-ready soil biodiversity indicators applicable to mountain ecosystems, which enable policy-makers to prioritise mountain soils for conservation (Guerra *et al*., 2021). In support of such developments, we reiterate previous calls (Maestre & Eisenhauer, 2019; Guerra *et al*., 2021, 2022) for improved monitoring of soil biodiversity and soil-related essential biodiversity variables and for increased efforts to identify hotspots of mountain soil biodiversity and endemism, as well as priority habitats in the light of ongoing and future global change. We also call for the systematic evaluation of the efficiency of protected areas in preserving mountain soil species and their functions (see e.g. Ciobanu *et al*., 2019). International initiatives such as SoilBON, the Global Soil Biodiversity Initiative, and the Global Soil Partnership of the United Nations Food and Agriculture Organisation represent effective opportunities for mountain soil scientists to engage with the endorsement of the Mountain Partnership (e.g. Stanchi *et al*., 2023), the Global Mountain Biodiversity Assessment, and other institutions committed to the conservation and sustainable use of mountain biodiversity. Besides political commitments and increased scientific efforts, awareness raising and education through effective communication methods (e.g. Steinwandter & Seeber, 2022) remain essential on our path to safeguarding sustainable mountain soils.

## VII. CONCLUSIONS

1. Despite a growing number of initiatives responding to the demand for data and knowledge on soil biodiversity, there are still major gaps and blind spots that exist, especially for the Global South and remote areas such as mountains.
2. This review intended to highlight the gaps in knowledge regarding mountains, which have become even more vulnerable due to ongoing land-use and climate change. Given the natural hazards and ecosystem services associated with mountain areas, maintaining soil health is of paramount importance. However, due to difficulties in data collection and the lack of a systematic assessment of the existing research corpus, addressing these challenges is proving challenging.
3. Here, we conducted a comprehensive review of the globally available data on biodiversity in temperate and continental alpine soils, above the treeline. This is, to our knowledge, the first time such data has been collated. Our systematic literature survey involved experts in the field of alpine soil biology, where we obtained an overview of the geographical distribution and number of studies focusing on alpine soil invertebrates, microbiota and cryptogams.
4. Our review has revealed research gaps in alpine regions outside of Europe and Central Asia, as well as for soil cryptogams and soil invertebrates, which have relatively limited data available in comparison to soil microbiota. Shortcomings were particularly notable among soil protists and soil invertebrates, and for the vast majority of uncultivated prokaryotes and fungi, for which functional or ecological descriptions are lacking. To address these issues, it will be necessary to improve geographic and taxonomic coverage. All these soil organisms have evolved over millennia to withstand the ever-growing extreme environmental conditions in mountain areas. Thus, there is a pressing need for a wider range of knowledge on all fronts.
5. We highlight three crucial areas for future research and policymaking on soil biodiversity in mountainous regions, emphasising their global distribution and the distinctive challenges posed by elevational gradients, remoteness and exposure to global change.
6. We call for a significant improvement in mountain soil biodiversity data, while we emphasise the need for enhancing the understanding of environmental drivers and consequences for biodiversity in mountain soils, advocating for improved spatial and temporal data resolution. Furthermore, we stress the importance of comparative analyses across different mountain ranges to inform conservation strategies in the face of global change.
7. Our review recommends clear political commitments, international collaboration, and the incorporation of biodiversity in mountain soils within global frameworks. This underscores the importance of raising awareness and providing education to promote the conservation and sustainable use of mountain soils.

## Supporting information

AppendixS1

## ACKNOWLEDGEMENTS

Michele Freppaz acknowledges the support of NBFC to University of Torino, funded by the Italian Ministry of University and Research, PNRR, Missione 4 Componente 2, ‘Dalla ricerca all’impresa’, Investimento 1.4, Project CN00000033. Andrea Britton and Andy Taylor were supported by the Scottish Government Rural and Environmental Science and Analytical Services (RESAS) as part of the Scottish Government Environment, Natural Resources and Agriculture (ENRA) Strategic Research Programme, project JHI-D4-3. Beat Frey acknowledges financial support from Fondation Petersberg pro Planta et Natura (grant number: 01.MZ/2023).

## VIII. SUPPORTING INFORMATION

Additional supporting information may be found online in the Supporting Information section at the end of the article.

**Appendix S1:** Detailed description of the methodology and statistical analyses, accompanied by additional tables and figures:

**Table S1.** List of search strings used to assess the available literature in ‘Web of Science’ focusing on alpine mountain soil biodiversity.

**Table S2.** Main steps of the semi-quantitative literature analysis to assess the number of available scientific papers focusing on alpine mountain soil biodiversity.

**Table S3.** List of the alpine regions and the encompassing mountain ranges used for this review.

**Table S4.** Number of scientific papers focusing primarily and secondarily on cryptogams in alpine mountain soils.

**Table S5.** Number of scientific papers focusing primarily and secondarily on microbiota (archaea, bacteria, fungi, and protists) in alpine mountain soils.

**Table S6.** Number of scientific papers focusing primarily and secondarily on fauna (macro-, meso-, and microfauna) in alpine mountain soils.

**Fig. S1.** Global map of scientific paper density per 1,000 km² on alpine soil biodiversity (cryptogams, soil microbiota and soil fauna) by mountain region.

**Fig. S2.** Comparison of the area of the eleven global alpine regions used in this review.

