## AppendixS1 for "Biodiversity in mountain soils above the treeline"

### I. MATERIAL AND METHODS

#### (1) Literature search

We performed a (semi-)quantitative literature analysis to assess the volume and geographical patterns of alpine mountain soil biodiversity research (including papers available until 11 November 2022). We chose 'Web of Science' (https://www.webofscience.com) as the source database because of its built-in functionalities to create complex searches using Boolean operators and a large number of search strings. We applied the search to the title, author, keyword, and abstract fields only. The Boolean search string consisted of five thematic areas, whose intersection defined the domain of knowledge of our interest, as described below.

**Mountain environments:** search terms that identify mountain environments generically, such as '*mountain**' and '*highland**'. We noticed – as also reported by Gurgiser *et al.,* (2022) – that many scientific studies do not necessarily include generic mountain descriptors in their abstracts and titles, therefore, we also included the specific names of mountain ranges (Snethlage *et al.*, 2022a) that (partially) overlap with the alpine bioclimatic belt (Testolin, Attorre & Jiménez-Alfaro, 2020), meaning zones above the natural treeline (i.e. alpine and nival zones).

**The alpine biome:** we limited the search outputs to studies (at least partially) conducted in the alpine bioclimatic belt, by including terms such as '*alpine*', '*treeline*', '*timberline*', '*glacier*' etc. and their most common alternative spellings (e.g. 'treeline' and 'tree line').

**The soil environment:** search terms to limit the results to the soil realm included '*soil**', '*edaph**', '*pedo**', '*subsurface*', etc.

**(Soil) biota**: names of relevant taxonomic groups were included to limit results to the major soil dwelling taxa such as cryptogams, bacteria, fungi, archaea, protists and invertebrates.

**Exclusion factors**: this group of keywords consisted of words to exclude tropical mountain ranges, as the focus of the review is on mountains in the temperate realm. It also included search terms to exclude archaeo- or paleo science.

The final list of keywords used is shown in Table S1, but for the list of mountain names see (Snethlage *et al.*, 2022a).

**Table S1.** List of search strings used to assess the available literature in 'Web of Science' focusing on alpine mountain soil biodiversity. The analysis includes literature that was available until 11 November 2022.

| |
|---|
| ‘alpine’ OR ‘subalpine’ OR ‘timber line*’ OR ‘timberline*’ OR ‘tree line*’ OR ‘treeline*’ OR ‘krummholz’ OR ‘cryogenesis’ OR ‘cryopedolog*’ OR ‘cryoturbat*’ OR ‘ice’ OR ‘glacial’ OR ‘glacier*’ OR ‘nival’ OR ‘firn’ OR ‘permanent snow’ OR ‘perennial snow*’ OR ‘perpetual snow’ OR ‘snow field’ OR ‘snow line’ OR ‘snowcap*’ OR ‘thermokarst’ OR ‘permafrost’ OR ‘gelisol*’ OR ‘cryosol*’ OR ‘cryophil*’ OR cryobiol* OR ‘psychrophil*’ OR ‘snow bed*’ OR ‘high* elevation*’ |
| AND |
| ‘aggregate stabilit*’ OR ‘albeluvisol*’ OR ‘alfisol*’ OR ‘andisol*’ OR ‘andosol*’ OR ‘anthrosol*’ OR ‘arenosol*’ OR ‘aridisol*’ OR ‘belowground’ OR ‘below-ground’ OR ‘calcisol*’ OR ‘cambisol*’ OR ‘cation exchange capacity’ OR ‘cryosol*’ OR ‘durisol*’ OR ‘edaph*’ OR ‘entisol*’ OR ‘epipedon*’ OR ‘fluvisol*’ OR ‘gelisol*’ OR ‘gleysol*’ OR ‘histosol*’ OR ‘humus’ OR ‘inceptisol*’ OR ‘leptosol*’ OR ‘mollisol*’ OR ‘myrmecosphere’ OR ‘pedodiversity’ OR ‘pedogenesis’ OR ‘pedogeograph*’ OR ‘pedolog*’ OR ‘pedon’ OR ‘pedosphere’ OR ‘planosol*’ OR ‘plant root*’ OR ‘podzol*’ OR ‘regosol*’ OR ‘rhizosphere*’ OR ‘rooting depth*’ OR ‘soil*’ OR ‘spodosol*’ OR ‘ultisol*’ OR ‘umbrisol*’ OR ‘epigeous’ OR ‘epigeal’ OR ‘epigean’ OR ‘epigeic’ OR ‘endogean’ OR ‘endogeic’ OR ‘soil crust’ OR ‘epigaeic’ |
| AND |
| ‘acari’ OR ‘acariform*’ OR ‘acidobacteri*’ OR ‘actinobacteri*’ OR ‘actinomycet*’ OR ‘agaricomycet*’ OR ‘algae’ OR ‘alphaproteobacter*’ OR ‘alveolat*’ OR ‘amoeboz*’ OR ‘Annelida’ OR ‘anthocerotophyt*’ OR ‘ants’ OR ‘Apterygota’ OR ‘arachnid*’ OR ‘araneae’ OR ‘archaea’ OR ‘archaeorhizomycet*’ OR ‘archaeplastid*’ OR ‘arthropod*’ OR ‘ascomycot*’ OR ‘bacillota’ OR ‘bacteri*’ OR ‘bacteroidet*’ OR ‘bacteroidia’ OR ‘basidiomycet*’ OR ‘Basidiomycetes’ OR ‘basidiomycot*’ OR ‘bathyarchaeot*’ OR ‘betaproteobacteri*’ OR ‘biocrust*’ OR ‘biological soil crust*’ OR ‘bryophyt*’ OR ‘bryophyte*’ OR ‘Bryopsida’ OR ‘carabid*’ OR ‘Caraboidea’ OR ‘centiped*’ OR ‘cetraria’ OR ‘cetrarioid ‘ OR ‘cetrarioid species’ OR ‘chasmophyt*’ OR ‘Chilognatha’ OR ‘chilopod*’ OR ‘choanozoa’ OR ‘chordeumatid*’ OR ‘chromadorea’ OR ‘chytridiomycot*’ OR ‘ciliat*’ OR ‘ciliate*’ OR ‘ciliophora’ OR ‘cladonia ‘ OR ‘Cladoniaceae’ OR ‘Clitellata’ OR ‘coleopter*’ OR ‘collembola*’ OR ‘Colpodea’ OR ‘conehead*’ OR ‘Crassiclitellata’ OR ‘crenaerchaeot*’ OR ‘crenarchaeot*’ OR ‘crustose lichen*’ OR ‘cryptobiotic surface crust’ OR ‘cryptogam*’ OR ‘cryptogamic crust*’ OR ‘cyanobacter*’ OR ‘Cystofilobasidiales’ OR ‘dictyosteli*’ OR ‘dictyostelid cellular slime mold*’ OR ‘dictyostelid cellular slime mould*’ OR ‘Dictyosteliida’ OR ‘diplopod*’ OR ‘dipluran*’ OR ‘Dipluridae’ OR ‘Diptera’ OR ‘DPANN’ OR ‘earthworm*’ OR ‘enchytraeid*’ OR ‘enoplea’ OR ‘Eupnoi’ OR ‘euryarchaeot*’ OR ‘firmicut*’ OR ‘flagellat*’ OR ‘formicid*’ OR ‘formicoid*’ OR ‘fruticose lichen*’ OR ‘fung*’ OR ‘gammaproteobacter*’ OR ‘gemmatimonadet*’ OR ‘Geophilidae’ OR ‘Geophilomorpha’ OR ‘glomerid*’ OR ‘glomeromycet*’ OR ‘glomeromycot*’ OR ‘gnaphosid*’ OR ‘grossglockneriid*’ OR ‘Halvaria’ OR ‘Harosa’ OR ‘hepatic*’ OR ‘Hexapoda’ OR ‘hornwort*’ OR ‘Hylocomiaceae’ OR ‘hymenopter*’ OR ‘hypha*’ OR ‘Hypnales’ OR ‘Icmadophilaceae’ OR ‘insect larva*’ OR ‘insect*’ OR ‘invertebrat*’ OR ‘isopod*’ OR ‘julid*’ OR ‘Lecanorales’ OR ‘Lecanoromycetes’ OR ‘leotiomycet*’ OR ‘leucosporidi*’ OR ‘lichen*’ OR ‘liverwort*’ OR ‘lumbricin’ OR ‘lycosid*’ OR ‘macroinvertebrate*’ OR ‘macrolichen*’ OR ‘Malacostraca’ OR ‘marchantiophyt*’ OR ‘mesoinvertebrate*’ OR ‘methanobacter*’ OR ‘micro organism*’ OR ‘microarthropod*’ OR ‘microb*’ OR ‘Microbotryomycetes’ OR ‘microeukaryote*’ OR ‘microinvertebrate*’ OR ‘microorganism*’ OR ‘milliped*’ OR ‘mite*’ OR ‘mollusc*’ OR ‘moss*’ OR ‘mosses’ OR ‘mrakia’ OR ‘mycelium’ OR ‘Mycetophilidae’ OR ‘mycetozoa’ OR ‘mycobiome*’ OR ‘mycorrhiz*’ OR ‘Myriapoda’ OR ‘naganishia’ OR ‘nanoarchaeot*’ OR ‘Negibacteria’ OR ‘nematocera’ OR ‘nematoceran larva*’ OR ‘nematod*’ OR ‘nitrososphaerot*’ OR ‘nitrosphaer*’ OR ‘oligochaet*’ OR ‘oniscidea’ OR ‘oomycet*’ OR ‘oomycot*’ OR ‘opilion*’ OR ‘oribatid mite*’ OR ‘Oribatida’ OR ‘parasitiform*’ OR ‘parmelia’ OR ‘patescibacter*’ OR ‘pauropod*’ OR ‘Pertusariales’ OR ‘PGPR’ OR ‘phalangiid*’ OR ‘Plant growth-promoting rhizobacteria’ OR ‘pleurocarp* moss’ OR ‘pleurozium ‘ OR ‘Posibacteria’ OR ‘potworm*’ OR ‘proteoarchaeot*’ OR ‘proteobacter*’ OR ‘proteobacteri*’ OR ‘protist*’ OR ‘protistan myxomycet*’ OR ‘protura*’ OR ‘Pseudofungi’ OR ‘pseudomonadota ‘ OR ‘pseudoscorpion*’ OR ‘rhizaria’ OR ‘rhizobacter*’ OR ‘rhodotorula’ OR ‘rotifer*’ OR ‘Sarcomastigota’ OR ‘Sarcoptiformes’ OR ‘scarabaeid*’ OR ‘Scarabaeoidea’ OR ‘slime mold*’ OR ‘slime mould*’ OR ‘soil animal*’ OR ‘soil arthropod*’ OR ‘soil biodiversit*’ OR ‘soil biological diversity’ OR ‘soil biota’ OR ‘soil crust community’ OR ‘soil fauna’ OR ‘soil invertebrat*’ OR ‘soil life’ OR ‘soil macrofauna’ OR ‘soil mesofauna’ OR ‘soil microbiome’ OR ‘soil microbiont’ OR ‘soil microbiota’ OR ‘soil microfauna’ OR ‘soil microflora’ OR ‘soil organism*’ OR ‘sordariomycet*’ OR ‘Sphagnaceae’ OR ‘Sphagnales’ OR ‘Sphagnopsida’ OR ‘sphagnum ‘ OR ‘spider*’ OR ‘Sporidiobolales’ OR ‘springtail*’ OR ‘staphylinid*’ OR ‘Staphylinoidea’ OR ‘Stelamoeba’ OR ‘stramenopil*’ OR ‘TACK’ OR ‘tardigrad*’ OR ‘terrabacteri’ OR ‘testate amoeba*’ OR ‘thamnolia ‘ OR ‘thaumarchaeot*’ OR ‘thermoleophil*’ OR ‘thermoplasmat*’ OR ‘thermoprot*’ OR ‘thomisid*’ OR ‘thysanoura*’ OR ‘thysanura*’ OR ‘Tremellaceae’ OR ‘Tremellales’ OR ‘Tremellomycetes’ OR ‘umbilicaria’ OR ‘verrucomicrob*’ OR ‘Vespoidea’ OR ‘woodlice’ OR ‘woodlouse’ OR ‘woseiarch*’ OR ‘yeast*’ OR ‘zygomycot*’ OR ‘cryophil*’ OR cryobiol* OR ‘psychrophil*’ OR ‘epigeous’ OR ‘epigeal’ OR ‘epigean’ OR ‘epigeic’ OR ‘endogean’ OR ‘endogeic’ OR ‘soil crust’ |

| |
|---|
| AND |
| *mountain** OR *cordillera** OR *alps* OR *alpine* OR *subalpine* OR *highland** OR *'high elevation*'* OR *'high altitude*'* |
| *+ selection of alpine high / mid latitude mountains from GMBA Mountain Inventory v2* |
| NOT |
| *tropical* OR *equatorial*<br>OR *'Angola*'* OR *'United Arab Emirates'* OR *'Antarctica'* OR *'Burundi'* OR *'Benin'* OR *'Burkina Faso'* OR *'Belize'* OR *'Brazil*'* OR *'Botswana'* OR *'Central African Republic'* OR *'Côte d'Ivoire'* OR *'Cameroon*'* OR *'Democratic Republic of the Congo'* OR *'Republic of Congo'* OR *'Colombia*'* OR *'Costa Rica*'* OR *'Cuba*'* OR *'Cyprus'* OR *'Dominica*'* OR *'Dominican Republic'* OR *'Ecuador*'* OR *'Egypt'* OR *'Eritrea*'* OR *'Ethiopia*'* OR *'Gabon'* OR *'Ghana'* OR *'Guatemala*'* OR *'French Guiana'* OR *'Guyana'* OR *'Honduras'* OR *'Haiti*'* OR *'Indonesia*'* OR *'Israel'* OR *'Jordan'* OR *'Kenya*'* OR *'Cambodia*'* OR *'Laos'* OR *'Lebanon'* OR *'Liberia'* OR *'Libya'* OR *'Sri Lanka*'* OR *'Madagascar'* OR *'Mali'* OR *'Mozambique'* OR *'Malaysia'* OR *'Namibia*'* OR *'New Caledonia'* OR *'Niger'* OR *'Nigeria'* OR *'Nicaragua'* OR *'Oman'* OR *'Panama'* OR *'Philippines'* OR *'Papua New Guinea'* OR *'Puerto Rico'* OR *'Reunion'* OR *'Rwanda'* OR *'Saudi Arabia'* OR *'Sudan'* OR *'Sierra Leone'* OR *'El Salvador'* OR *'Somalia'* OR *'South Sudan'* OR *'Suriname'* OR *'Syria'* OR *'Chad'* OR *'Togo'* OR *'Thailand'* OR *'Tanzania*'* OR *'Uganda*'* OR *'Uruguay'* OR *'Venezuela*'* OR *'Vietnam'* OR *'Yemen'* OR *'Zambia'* OR *'Zimbabwe'*<br>OR *fossil** OR *jurassic* OR *miocene* OR *neogene* OR *palaeo** OR *paleo** OR *palyno** OR *permian* OR *quaternary* OR *Pleistocene* OR *Triassic*<br>OR *'lake'* OR *(forest** NOT *(meadow** OR *grassland** OR *tundra* OR *pasture** OR *heath* OR *peat* OR *bog* OR *fen* OR *scree))* |

### (2) Data processing

The paper selection process consisted of three steps starting with a 'naïve search' (Grames *et al.*, 2019) in 'Web of Science': lead authors provided keywords (mainly key taxa names) for the nine main organismic groups considered in the review and some keywords for the other alpine thematic areas. From the results of this 'naïve search', we extracted further keywords with the R package LITSEARCHR (Grames *et al.*, 2019) in the open-source statistical programming environment R (version 4.3.1, R Core Team, 2023) and with the Global Names Finder (Mozzherin, Myltsev & Zalavadiya, 2023; https://gnrd.globalnames.org/). We reviewed and cleaned these additional keywords and taxa lists and added them to the revised search string making sure to include common spelling alternatives (e.g. hyphenated variants: not only microarthropods but also micro-arthropods). The 'Web of Science' search resulted in 1,997 potentially relevant publications (see Table S2). To these, the authors added an additional 225 references from their personal reference databases.

We then associated each paper with the taxa (i.e. species, genus, family etc.) extracted from the corpus with the Global Names Finder app with the STRINGR R package (Wickham, 2022) and linked them to their corresponding organismic group(s). For example, we attributed each occurrence of the taxon 'spider' in a paper to the organismic group 'macro-invertebrates'. We then shared the publications lists grouped by organismic group with the corresponding lead authors for a round of validation, which resulted in the elimination of 680 references. This brought the final number of publications to 1,557. For georeferencing, also using the R package STRINGR, we extracted mountain range names (Snethlage *et al.*, 2022a) and country names from each publication and added 27 mountain range names manually. We then aggregated 810 paper–mountain range associations to the 11 larger alpine regions using the hierarchical structure of the 'GMBA Mountain Inventory v2'. This allowed us to extract statistics on the number of references by organismic group and alpine mountain region. Further, we calculated the area of each alpine mountain region; see Table 1 in the main text, and Table S3 and Fig. S2 to be able to calculate the number of papers per unit area of alpine habitat.

**Table S2.** Main steps of the semi-quantitative literature analysis to assess the number of available scientific papers focusing on biodiversity in mountain soils above the treeline. These results correspond to a comprehensive literature research made in 'Web of Science' (i.e. available papers until 11 November 2022).

| Phase of the semi-quantitative literature review | # papers |
|---|---|
| 'Web of Science' search | 1,997 |
| Added by the authors | 240 |
| Eliminated after review and validation by the authors | 680 |
| Kept for analysis | 1,557 |
| GMBA mountain ranges extracted | 810 |
| Mountain range names added manually | 27 |
| Country name extracted | 690 |
| Target* organismic group extracted AND georeferenced (mountain or country) | 972 |
| **Primary target* organismic group extracted AND georeferenced (mountain or country)** | **708** |

* target organismic group: organismic groups discussed in this paper, others being plants, vertebrates etc.

As many papers about biodiversity in mountain soils covered more than one organismic group, we counted the number of taxon mentions per organismic group in each paper's title, abstract and keywords, and attributed the flag 'primary focus' of the paper to the organismic group with the most taxa, and 'secondary focus' to the one with the second highest number of taxa. For example, the flag primary focus would be 'macro-invertebrates', and secondary focus 'fungi' for a paper with the extracted strings 'earthworm', 'millipedes', 'arthropoda' (three macro-invertebrate groups), and 'agaricomycetes' (a group of fungi).

**Table S3.** List of the eleven alpine regions and the encompassing mountain ranges used for this review. Main ranges are ordered alphabetically; subranges that are listed in the text are in parentheses in italic. The mountain range names are taken from the 'GMBA Mountain Inventory v2' (see Snethlage *et al.*, 2022b).

| Nr. | Alpine region | Mountain ranges |
|---|---|---|
| 1 | North American Cordillera | Alaska Range – Brooks Range – Alaska Intermountain Ranges – Yukon Intermountain Ranges – Mackenzie Mountains – South-Central Alaska – Saint Elias Mountains (*Southern Icefield Ranges*) – Coast Mountains – British Columbia Interior (*Chilcotin Plateau*) – Far Northern Rockies – Canadian Rockies – Columbia Mountains – Cascade Range – Central Montana Rocky Mountains – Idaho-Bitterroot Rocky Mountains – Columbia Plateau – Oregon Coast Range – Northern California Coast Ranges – Sierra Nevada – Western Rocky Mountains – Southern Rocky Mountains – Colorado Plateau – Greater Yellowstone Rockies – Western Rocky Mountains |
| 2 | Appalachians & Northeast Ranges | Adirondack Mountains – Appalachian Mountains (*Great Smoky Mountains*) – Arctic Cordillera – Central Labrador Ranges – Laurentian Mountains, |
| 3 | Northern Europe | British Isles – Fell Lapland – Northern Scandes – Southern Scandes (*Vestvidda*) |
| 4 | North Asia | Anadyr Highlands – Aldan Mountains – Bureya Region – Central Siberian Plateau – Chersky Range – Chukotka Mountains – Dzhugdzhur Mountains – Kamchatka Peninsula – Kolyma Mountains – Koryak Mountains – Kuznetsk Alatau – Lena-Angara Plateau – Mongolian Altai – Mongolian Highlands – Northern Altai – Northern Baikal Mountains – Patom Highlands – Sakhalin Peninsula – Sikhote-Alin – Stanovoy Highlands – Stanovoy Range – Ural Mountains – Verkhoyansk Range – Yana-Oymyakon Highlands – Western Sayan – Yankan - Tukuringra - Soktakhan - Dzhagdy group of mountain ranges – Yukaghir Highlands |
| 5 | Central & Southern Europe | Bohemian Massif (*Eastern Sudetes*) – Carpathian Mountains – Dinaric Alps – European Alps (*High Tauern, Ötztal Alps, Uri Alps*) – Hellenides – Pyrenees – Rila-Rhodope Massif |
| 6 | Caucasus | Alborz Mountains – Armenian Highlands – Caucasus Mountains – Pontic Mountains – Taurus Mountains |
| 7 | Central Asia | Alashan Plateau – Altyn-Tagh – Balochistan Ranges – Bayan Har Mountains – Hengduan Shan – Himalaya (*Zanskar Range*) – Hindu Kush – Karakoram – Kunlun Mountains – Min Mountains – Pamir Mountains – Qiangtang – Qilian Mountains – Qionglai Shan – Tanggula Mountains – Tian Shan – Transhimalaya |
| 8 | East Asia | Changbai Mountains – Hokkaido – Honshu (*Kantō Mountains*) – Korean Peninsula |
| 9 | Andes & South America | Cordillera Central – Cordillera de la Costa – Cordillera Occidental – Cordillera Oriental (*Cordillera de Vilcanota*) – Dry Andes – Meseta Patagónica – Patagonian Andes (*Cordillera Darwin*) – Sierras Pampeanas |
| 10 | Afro-alpine Region | Albertine Rift Mountains – Eastern Rift mountains – Ethiopian Highlands – Drakensberg |
| 11 | Australia & New Zealand | Fiordland – North Island – Northwest Ranges – Southern Alps – Southern Great Dividing Range – Tasmania |

### II. RESULTS

The result of the literature search per organismic group (cryptogams, soil microbiota and invertebrates) is illustrated in Fig. 1 in the main text. The absolute numbers of publications per alpine mountain area and soil organism group are given in the Tables S4–S6. Given the large differences in geographic extent of the examined alpine mountain regions (see Table 1 in main text), we normalized the number of publications for each alpine area, resulting in references-per-km² densities (Fig. S1).

In Central Asia, particularly in the Himalaya and Tibetan Plateau, the publication output is significantly lower when corrected for the area of the alpine habitat (Fig. 1 and Fig. S1). Both absolute and relative paper output remains high in Central & Southern Europe, primarily driven by a high output of papers focused on research conducted in the European Alps). East Asia is the second focal point for biodiversity research in mountain soils above the treeline highlighted by numerous publications centred on the Changbai Mountains (East Asia). The disproportionately high relative paper output for the Afro-alpine region is likely driven by the limited expanse of alpine habitat in that area.

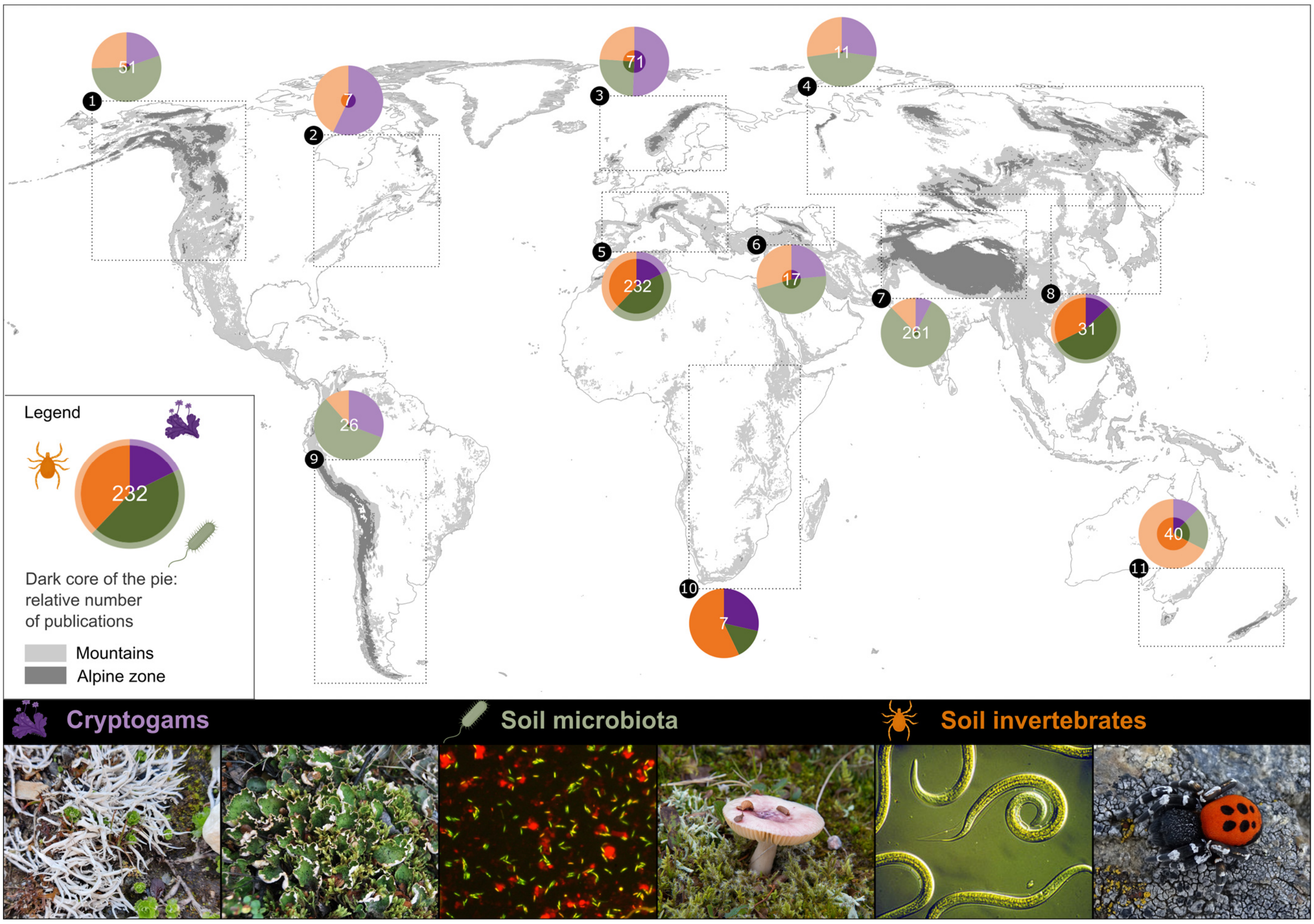

**Fig. S1.** Global map of scientific paper density per 1,000 km² on biodiversity (cryptogams, soil microbiota and soil invertebrates) in mountain soils above the treeline by mountain region. The dark core of the pies represents the number of publications in the respective area compared to the number of the region with the highest paper density (i.e. Afro-alpine Region, highlighted). The regions were named here as (1) North American Cordillera, (2) Appalachians & Northeast Ranges, (3) Northern Europe, (4) North Asia, (5) Central & Southern Europe, (6) Caucasus, (7) Central Asia, (8) East Asia, (9) Andes & South America, (10) Afro-alpine Region, and (11) Australia & New Zealand. Photos from left to right: Cryptogams: Arctic-alpine lichen *Thamnolia vermicularis*, arctic-alpine lichen *Peltigera aphthosa* (credit: Bettina Weber); Soil microbiota: DNA stained microscope preparation of soil bacteria (credit: Nadine Praeg & Paul Illmer), *Russula* sp. (credit: Andrea J. Britton); Soil fauna: Nematodes and a male velvet spider *Eresus sandaliatus* (credits: CSIRO Entomology and Michael Steinwandter, respectively).

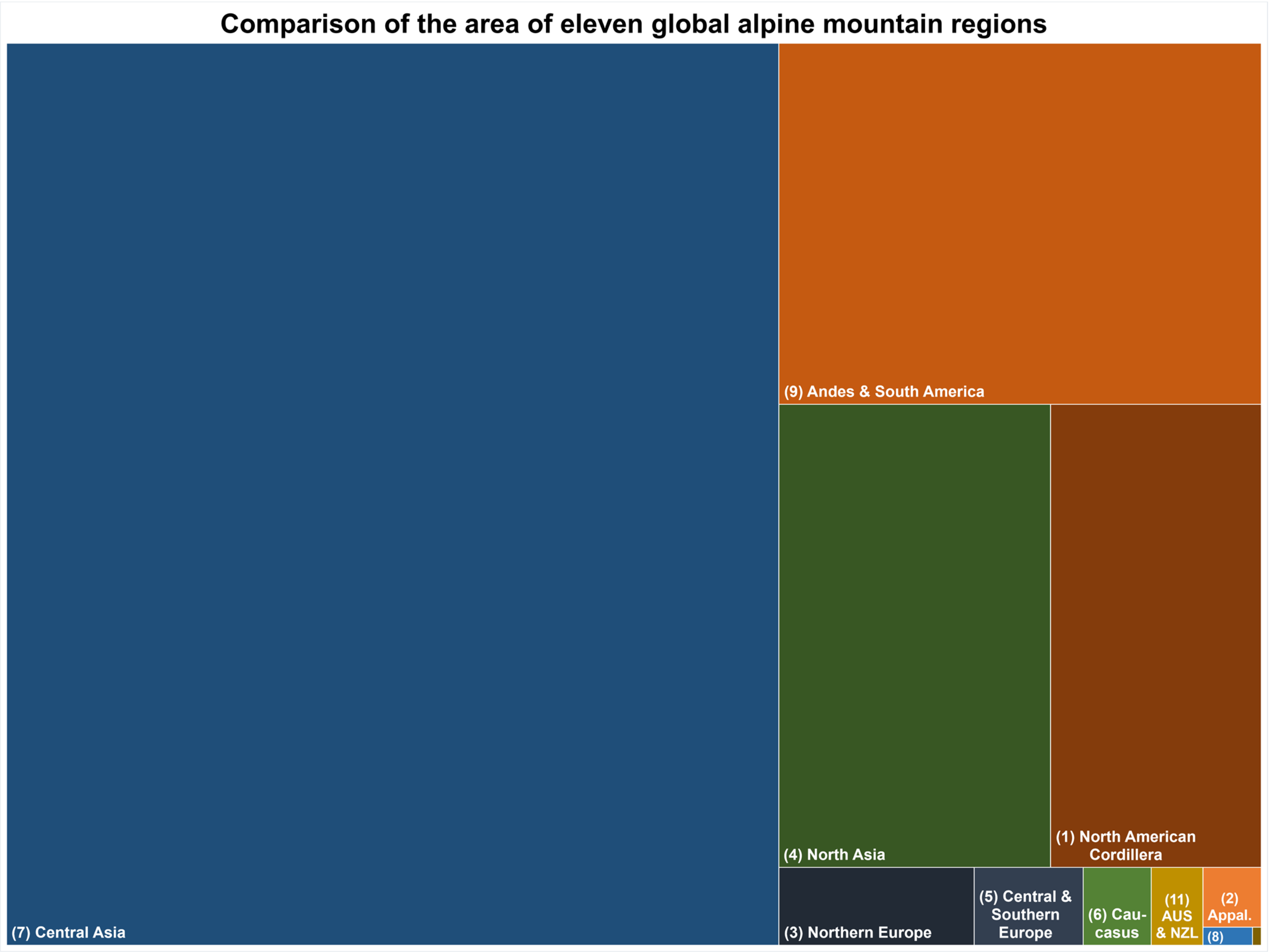


**Fig. S2.** Comparison of the area [km$^2$] of the eleven global alpine regions considered in this review. Size values for each alpine region are detailed in Table 1 (main text). Names of relatively small alpine areas are indicated: (2) Appalachians & Northeast Ranges, (8) East Asia; (11) Australia & New Zealand; the unlabelled rectangle in the lower right corner represents the smallest alpine region, the Afro-alpine Region.

**Table S4.** Number of scientific papers per alpine region focusing primarily and secondarily on cryptogam diversity in mountain soils above the treeline. These results correspond to a comprehensive literature research made in 'Web of Science' (i.e. available papers until 11 November 2022). Others refer to cryptogams not categorized as biocrusts (e.g. lichens).

| **Nr.** | **Alpine region** | **Number of scientific articles** | | | | | |
|---|---|---|---|---|---|---|---|
| | | **Biocrusts** | **Others** | ***Sum of first*** | **Biocrusts** | **Others** | ***Sum of second*** |
| 1 | North American Cordillera | – | 10 | *10* | 2 | 6 | *8* |
| 2 | Appalachians & Northeast Ranges | – | 4 | *4* | – | – | *0* |
| 3 | Northern Europe | 1 | 35 | *36* | 6 | 14 | *20* |
| 4 | North Asia | – | 3 | *3* | 0 | 1 | *1* |
| 5 | Central & Southern Europe | 7 | 34 | *41* | 15 | 43 | *58* |
| 6 | Caucasus | – | 4 | *4* | – | 7 | *7* |
| 7 | Central Asia | 7 | 13 | *20* | 10 | 33 | *43* |
| 8 | East Asia | – | 4 | *4* | 1 | 6 | *7* |
| 9 | Andes & South America | – | 8 | *8* | 4 | 7 | *11* |
| 10 | Afro-alpine Region | 1 | 1 | *2* | 1 | 1 | *2* |
| 11 | Australia & New Zealand | – | 5 | *5* | 2 | 8 | *10* |
| | **Sums** | **16** | **121** | ***137*** | **41** | **126** | ***167*** |

**Table S5.** Number of scientific papers per alpine region focusing primarily and secondarily on microbial diversity (archaea, bacteria, fungi, and protists) in mountain soils above the treeline. These results correspond to a comprehensive literature research made in 'Web of Science' (i.e. available papers until 11 November 2022).

| **Nr.** | **Alpine region** | **Number of scientific articles** | | | | | | | | | |
|---|---|---|---|---|---|---|---|---|---|---|---|
| | | **Archaea** | **Bacteria** | **Fungi** | **Protists** | ***Sum of first*** | **Archaea** | **Bacteria** | **Fungi** | **Protists** | ***Sum of second*** |
| 1 | North American Cordillera | 1 | 6 | 20 | 1 | *28* | – | 9 | 10 | 3 | *22* |
| 2 | Appalachians & Northeast Ranges | – | – | – | – | – | – | – | 1 | – | *1* |
| 3 | Northern Europe | – | 4 | 13 | 1 | *18* | – | 9 | 18 | 1 | *28* |
| 4 | North Asia | – | 1 | 4 | – | *5* | – | 1 | – | – | *1* |
| 5 | Central & Southern Europe | 5 | 33 | 56 | 9 | *103* | 11 | 57 | 66 | 16 | *150* |
| 6 | Caucasus | – | 2 | 4 | 2 | *8* | – | 3 | 4 | 2 | *9* |
| 7 | Central Asia | 8 | 129 | 68 | 4 | *209* | 29 | 127 | 113 | 10 | *279* |
| 8 | East Asia | 1 | 7 | 8 | 1 | *17* | – | 16 | 12 | 2 | *30* |
| 9 | Andes & South America | – | 7 | 5 | 3 | *15* | – | 7 | 9 | 2 | *18* |
| 10 | Afro-alpine Region | – | – | – | 1 | *1* | – | 1 | – | – | *1* |
| 11 | Australia & New Zealand | – | 1 | 6 | 1 | *8* | – | 2 | 4 | 2 | *8* |
| | **Sums** | **15** | **190** | **184** | **23** | ***412*** | **40** | **232** | **237** | **38** | ***547*** |

**Table S6.** Number of scientific papers per alpine region focusing primarily and secondarily on invertebrate diversity (macro-, meso-, and micro-invertebrates) in mountain soils above the treeline. These results correspond to a comprehensive literature research made in 'Web of Science' (i.e. available papers until 11 November 2022).

| **Nr.** | **Alpine region** | **Number of scientific articles** | | | | | | | |
|---|---|---|---|---|---|---|---|---|---|
| | | **Macro-inv.** | **Meso-inv.** | **Micro-inv.** | ***Sum of first*** | **Macro-inv.** | **Meso-inv.** | **Micro-inv.** | ***Sum of second*** |
| 1 | North American Cordillera | 9 | 2 | 2 | *13* | 1 | 1 | 1 | *3* |
| 2 | Appalachians & Northeast Ranges | 1 | – | 2 | *3* | 1 | 1 | – | *2* |
| 3 | Northern Europe | 9 | 5 | 3 | *17* | 7 | 5 | 1 | *13* |
| 4 | North Asia | 1 | 2 | – | *3* | 1 | 1 | – | *2* |
| 5 | Central & Southern Europe | 62 | 18 | 8 | *88* | 19 | 13 | 8 | *40* |
| 6 | Caucasus | 4 | 1 | – | *5* | 4 | 6 | 2 | *12* |
| 7 | Central Asia | 14 | 2 | 16 | *32* | 5 | 3 | 4 | *12* |
| 8 | East Asia | 4 | 4 | 2 | *10* | 4 | – | – | *4* |
| 9 | Andes & South America | 3 | – | – | *3* | 2 | 1 | 1 | *4* |
| 10 | Afro-alpine Region | 3 | – | 1 | *4* | 0 | 3 | – | *3* |
| 11 | Australia & New Zealand | 8 | 18 | 1 | *27* | 5 | 3 | – | *8* |
| | **Sums** | **118** | **52** | **35** | ***205*** | **49** | **37** | **17** | ***103*** |
